# Comparison of six methods for stabilizing metapopulation dynamics and the robustness of performance rankings to census noise

**DOI:** 10.1101/2024.02.04.578847

**Authors:** Akanksha Singh, Sudipta Tung

**Affiliations:** Indian Institute of Science Education and Research, Mohali, Punjab, India-140306; Integrated Genetics and Evolution Laboratory (IGEL), Department of Biology, Ashoka University, Sonipat, Haryana, India, 131029; Max Planck Institute for Evolutionary Biology, Plön, Germany, 24306; Integrated Genetics and Evolution Laboratory (IGEL), Evolutionary and Organismal Biology Unit (EOBU), Jawaharlal Nehru Centre for Advanced Scientific Research (JNCASR), Bangalore, Karnataka, India, 560064

**Keywords:** Constancy, effective population size, effort of implementation, persistence, population stability, spatially-structured populations

## Abstract

Stabilizing extinction-prone populations is a central challenge in conservation, particularly for species with overcompensatory dynamics that generate boom–bust cycles. In spatially structured populations, dispersal can synchronize these fluctuations and amplify instability, while imperfect census data can misguide interventions, making both factors critical determinants of management outcomes. Using a two-patch metapopulation model based on the Ricker equation, we evaluate six rule-based population control methods under spatial coupling and census uncertainty. We compare their relative performance across 12 ecological conditions defined by two intrinsic growth rates, two carrying capacities, and three dispersal scenarios, using constancy-targeted and persistence-targeted analyses that integrate multiple stability metrics with implementation cost. Uncontrolled metapopulations exhibited distinct stability profiles across parameter combinations. Against these baseline regimes, augmenting populations to a fixed fraction of their previous size (Adaptive Limiter Control) emerged as the strongest overall performer under persistence-targeted analyses, while augmenting populations whenever they fell below a threshold (Lower Limiter Control) performed most favourably overall under constancy-targeted analyses. Under simulated census error, the relative ranking of methods was often broadly preserved across noise levels. Our results provide a comparative framework for evaluating rule-based interventions in spatially structured populations exhibiting overcompensatory (Ricker-type) dynamics, including semelparous species with high fecundity.

## 1. Introduction

Populations rarely exist in stable equilibrium. Complex temporal fluctuations—periods of explosive growth followed by sharp crashes, or oscillations that persist for years—are ubiquitous in natural populations and represent a primary driver of extinction risk (Lundberg et al. 2000; Clark and Luis 2020). For conservation biologists and managers, such erratic dynamics pose a fundamental challenge: persistent large fluctuations increase mean extinction frequency, episodic low population sizes erode genetic diversity, and unpredictability complicates the allocation of limited management resources (Escudero et al. 2004; Smith and Meerson 2016). The success of any conservation intervention is therefore judged not merely by whether populations persist (persistence stability), but by whether fluctuations remain small (constancy stability), whether effective population size remains high to maintain genetic diversity, and whether the cost of intervention remains reasonable (Tung et al. 2014). These multiple, sometimes conflicting objectives— fluctuation amplitude, mean extinction frequency, effective population size, and cost—form the core metrics by which conservation managers evaluate whether a particular strategy has succeeded.

Over the past three decades, theoretical ecologists have proposed several rule-based control methods designed to stabilize inherently unstable populations by systematically adding or removing individuals according to simple rules based on current population size (McCallum 1992; Corron et al. 2000; Hilker and Westerho 2005; Dattani et al. 2011; Sah et al. 2013; Tung et al. 2014). The practical appeal of such methods is substantial: they require minimal knowledge of underlying system parameters and can be deployed immediately in real populations without detailed characterization of demographic rates. For example, Constant Pinning (CP) adds a fixed number of individuals at each census interval, shifting recruitment curves upward to convert chaotic dynamics into periodic or stable states (McCallum 1992; Parthasarathy and Sinha 1995; Solé et al. 1999). Lower Limiter Control (LLC) restrains the cost of augmentation by adding individuals only when population size drops below a predetermined threshold (Hilker and Westerho 2005), while Upper Limiter Control (ULC) achieves the opposite effect through strategic culling whenever populations exceed an upper threshold (Hilker and Westerho 2005). Both Limiter Control (BLC) combines restocking and culling to maintain populations within a target range (Tung et al. 2014). Adaptive Limiter Control (ALC) scales the number of added individuals as a proportion of the previous generation’s population size, creating density-dependent feedback that traps dynamics around carrying capacity (Sah et al. 2013). Target-Oriented Control (TOC) guides populations toward specific targets by adding or removing individuals proportional to deviation from the target population size (Dattani et al. 2011).

Despite representing a diverse spectrum of practical strategies with distinct decision rules and cost–benefit profiles, existing population control methods differ markedly in their trade-offs among key management targets, making it non-trivial to identify which method performs best under given ecological conditions. To address this challenge, a previous study (Tung et al. 2014) conducted the first comprehensive comparison of the above mentioned six methods using the Ricker population growth model within a unified theoretical framework. These methods were selected because they are implementable in empirical systems (Gusset et al. 2009), theoretically well-established (McCallum 1992; Solé et al. 1999; Hilker and Westerho 2005; Dattani et al. 2011; Tung et al. 2014), and empirically validated (Dey and Joshi 2013; Sah et al. 2013; Tung et al. 2016a,b). Together, they capture the main dimensions of population intervention, i.e. magnitude, direction (addition versus removal), and adaptiveness, and constitute a representative subset of theoretically proposed strategies. The Ricker model used in this study captures overcompensatory density dependence and boom-bust cycles characteristic of many biological populations including semelparous species with high intrinsic growth rates. It can be derived from first principles for randomly distributed individuals undergoing scramble competition (Brännström and Sumpter 2005) and has successfully predicted dynamics across diverse taxa, including bacteria, fungi, ciliates, insects, and fish (Ricker 1954; Sheeba and Joshi 1998; Denney et al. 2002; Ives et al. 2004; Fryxell et al. 2005; Ponciano et al. 2005; Dey and Joshi 2006), making it an ideal foundation for generalizable predictions. Tung et al. (2014) evaluated the control methods’ performance across four combinations of intrinsic growth rate (*r*) and carrying capacity (*K*) using composite scores integrating mean extinction frequency, fluctuation magnitude, effective population size, and cost. Their key findings were: methods involving culling (ULC, BLC, TOC) improved persistence (the probability of long-term population survival) more than constancy (the reduction of fluctuation magnitude), whereas restocking-only methods (CP, LLC, ALC) showed the opposite pattern. Critically, no single method dominated across all targets or conditions—relative efficacy depended strongly on *r* and *K*. Overall, Lower Limiter Control emerged as the most favourable strategy, achieving constancy while maintaining high effective population size and low cost under most single-population conditions.

However, this important study involved only spatially isolated populations. But natural populations rarely exist in isolation; they are more often part of spatially structured networks of subpopulations connected by dispersal, collectively termed metapopulations (Gyllenberg et al. 1993; Hanski 1998). Within such systems, dispersal can have both stabilising and destabilising effects: it may reduce local crowding and rescue declining subpopulations, but it can also synchronise fluctuations across patches, promoting simultaneous crashes (Stacey et al. 1997; Dey and Joshi 2006). Therefore, dispersal is a major determinant of stability in spatially structured populations, and its opposing effects imply that the relative performance of control methods is unlikely to transfer uniformly from isolated populations to spatially connected ones. Under high dispersal, we expect that increasing synchrony will make reducing overshoots more important, thereby giving culling-based methods (ULC, BLC, TOC) a relative advantage. Conversely, under low or asymmetric dispersal, we expect that preventing local crashes will become more important, leading augmentation-based methods (CP, LLC, ALC) to perform more favourably. We also expect methods incorporating density-dependent feedback (ALC, TOC) to be more robust to dispersal-induced shifts in local densities than fixed-rule approaches (CP, LLC, ULC), because control rules are applied separately to each patch and proportional adjustments can therefore accommodate patch-specific differences in baseline population size. Consequently, the very processes that shape dynamics in metapopulations can cause control methods to perform differently than they do in isolated populations, making single-population insights an unreliable guide for predicting metapopulation outcomes. While metapopulation-scale control has received attention in applied settings such as invasive-species management (e.g. Bogich and Shea 2008; Walker et al. 2015; Mallela and Hastings 2022; Kallis et al. 2023), systematic evaluation of rule-based methods for stabilizing extinction-prone metapopulations remains uncommon, with most previous work centred on single isolated populations or only a narrow range of spatial conditions (Dey and Joshi 2006; Sah et al. 2013).

This knowledge gap is compounded by a second, equally important problem: all rule-based control methods rely fundamentally on census counts, yet real censuses are inherently imperfect. Census data suffer from observation error, observer bias, and misidentification (Miller et al. 2011; Brack et al. 2018). Modern monitoring technologies do not eliminate these errors, instead they merely shift their character (Brack et al. 2018). Remarkably, no theoretical or empirical study has systematically examined how different control methods respond to census uncertainty in any population context, let alone in spatially structured systems. A commentary (Milner-Gulland and Shea 2017) on this topic documents how observation uncertainty can render management ineffective or even counterproductive, yet such uncertainty is routinely absent from theoretical comparisons of control strategies. When managers apply control methods tuned to particular *r-K* combinations under the assumption of perfect information, but must rely on imperfect census data in practice, do different magnitudes and forms of monitoring error (white noise, systematic over- or underestimation) alter the relative ranking of control methods? Does migration magnitude and symmetry affect the robustness of performance rankings of methods and thereby change which methods are preferred? These questions remain unanswered.

Here, we address both gaps by evaluating the same six rule-based control methods within a two-patch metapopulation framework, using an identical theoretical model, parameter space, and performance metrics as the single-population study (Tung et al. 2014). We deliberately preserve this modelling consistency to enable qualitative comparison with previously reported single-population results, providing a synthesis of control performance across spatial scales. We first characterize unperturbed metapopulation dynamics across symmetric and asymmetric dispersal regimes, quantifying how constancy, persistence, effective population size, and synchrony vary across a range of *r-K* combinations and migration rates. This baseline establishes how the interplay between intrinsic growth rate, carrying capacity, and dispersal shapes metapopulation stability in the absence of intervention. We then evaluate how each control method performs across the same *r-K* combinations and using the same Composite Performance Score (CPS) proposed in the previous study (Tung et al. 2014) under both symmetric and asymmetric migration. Finally, we test the robustness of method rankings to three types of census noise, white noise, positive bias (overestimation), and negative bias (underestimation), reflecting the diversity of real-world monitoring challenges.

We also qualitatively compare metapopulation outcomes with the corresponding single-population results and find that spatial structuring does not fundamentally alter the relative ranking of control methods. Our results reveal that Lower Limiter Control consistently delivers target constancy while maintaining high effective population size, high persistence probability, and low implementation cost across metapopulation conditions. When control parameters are instead tuned to achieve target persistence, Adaptive Limiter Control emerges as the most effective method, delivering favourable combinations of high effective population size, low mean extinction frequency, and low cost. Strikingly, when faced with census errors, the relative ranking of methods remains largely unchanged, suggesting that imperfect information is unlikely to substantially alter the choice of which control method to use. By integrating insights from both single-population and metapopulation studies into a unified comparative framework, and by evaluating how each method performs under realistic levels of census uncertainty, this work provides a clearer, evidence-based foundation for selecting stabilization strategies in spatially structured populations, particularly under the data-limited conditions common in ecological management.

## 2. Materials and Methods

### 2.1. Metapopulation framework and population growth model

We simulated a two-patch metapopulation under homogeneous environmental conditions. Population dynamics within each patch followed the Ricker model (Ricker 1954):

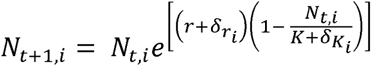

where *r* is the baseline intrinsic growth rate, *K* the baseline carrying capacity of each patch, and *N_t,i_* the population size in patch *i* at generation *t*. Thus, the two patches share the same baseline *r* and *K*. The Ricker formulation captures overcompensatory, discrete-time dynamics arising from scramble competition, in which density-dependent mortality intensifies sharply at high densities and produces boom–bust cycles characteristic of organisms with high fecundity and discrete, non-overlapping generations (May and Oster 1976). To incorporate demographic and environmental variability, patch-specific stochastic noise in *r* and *K*, δ*_ri_* and δ*_Ki_* respectively, were drawn independently at each time step from uniform distributions U[−0.2, 0.2] and U[−0.1K, 0.1K], respectively. These stochasticity levels follow the single-population study (Tung et al. 2014) and are included to introduce biologically plausible variation and prevent deterministic trajectories from remaining identical when initiated from the same starting population size. The noise amplitudes are sufficiently small that they do not alter the qualitative dynamical regime of the Ricker model or directly induce extinction (e.g., by driving *r* or *K* to unrealistically low values). Consistent with this, extinctions remain rare in the low-*r* scenario, indicating that the noise itself does not push populations below the extinction threshold.

Because the Ricker update produces continuous values while *N_t,i_* represents discrete individuals, we treated all population sizes as integer-valued state variables. Accordingly, Ricker outputs were rounded to the nearest integer to reflect demographic discreteness and to capture the lattice effects known to influence extinction dynamics (Henson et al. 2001; Domokos and Scheuring 2004). When population size fell below four individuals, a 50% probability of stochastic extinction was imposed to mimic demographic accidents, mating failure, or skewed sex ratios that result in no viable offspring. These choices replicate the single-population framework (Tung et al. 2014) to enable qualitative comparison. Together, integerization and the stochastic-extinction rule break the scale invariance of the Ricker map and make carrying capacity (*K*) meaningfully affect extinction risk: low-*K* populations are more vulnerable because coarser rounding relative to population size increases the likelihood of falling below the extinction threshold.

Following local population updates, a fixed proportion (*m_i_*) of individuals emigrated from patch *i* to the other, with the number of emigrants also rounded to the nearest integer, representing symmetric or asymmetric dispersal between subpopulations. The post-dispersal population size (*b_t+1_*, *_i_*) served as the census size for applying population control methods in the *i*th patch, while the post-control size (*a_t+1_*, *_i_*) acted as the input for the next Ricker iteration. Under unperturbed dynamics, *b_t+1, i_ =a_t+1, i_*. A schematic of the metapopulation structure and event sequence is shown in Figure 1.

**Figure 1.**
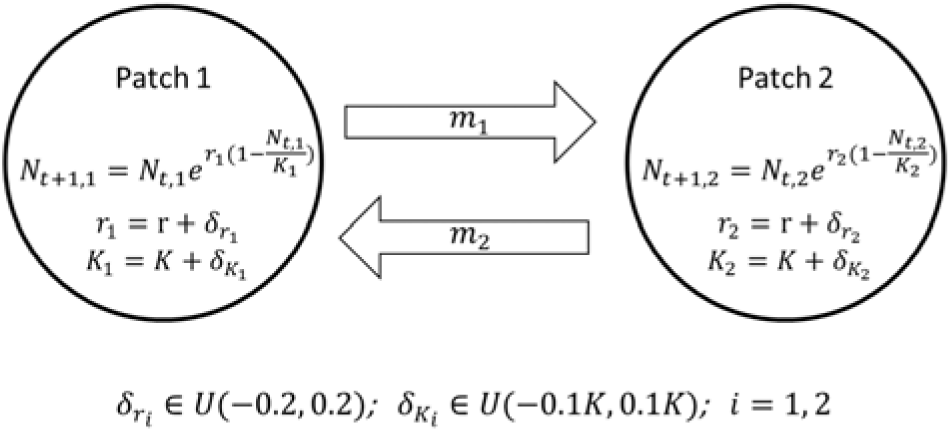
Simulation schematics. The schematic diagram shows the two-patch metapopulation system used for this study, where local dynamics each patch is modelled using Ricker growth model. We incorporated noise in both intrinsic growth rate (*r*) and carrying capacity (*K*) parameters in each iteration of the simulations by drawing random numbers from a uniform distribution (indicated by *U*) in the figure.

### 2.2. Baseline Metapopulation Dynamics: Parameter Space and Dispersal Regimes

To characterize unperturbed metapopulation dynamics, we explored a broad parameter space of intrinsic growth rate (*r*) and carrying capacity (*K*). The parameter ranges—*r* from 2.7 to 4.5 (step size 0.1) and *K* from 20 to 350 (step size 5)—encompass and extend the factorial combinations of *r* (2.8, 4.0) and *K* (60, 300) used subsequently for comparing the performance of the population control methods (see section 2.6). Lower *r* values were excluded since chaotic dynamics in the Ricker model only emerge above *r* ≈ 2.69, representing the onset of complex, extinction-prone regimes where control interventions are most relevant. Below this threshold, dynamics are monotonic or damped and inherently stable, offering little opportunity to meaningfully compare control strategies.

Simulations were performed under two symmetric dispersal scenarios with low (*m = m = 0.1*) and high (*m = m = 0.3*) migration rates, following empirically informed ranges from a previous study (Dey and Joshi 2006). To assess the effects of asymmetric dispersal, which reflects directional migration and unequal emigration tendencies between subpopulations, we further varied *m* and *m* independently from 0.01 to 0.49 in increments of 0.02 for two contrasting ecological regimes – (*r*, *K*) = (2.8, 60) and (4.0, 300), noting that migration values near 0.5 correspond to near-complete mixing between patches.

### 2.3. Stability Metrics

We quantified metapopulation stability using four complementary metrics:

**Constancy**: Temporal variability in metapopulation size was measured using the *Fluctuation Index* (*FI*) = Σ|*N_t+1_ − N_t_*| / (N × *T*), where the sum is taken over *t* = 0 to *T−1*, N is mean metapopulation size and *T* is the number of generations (Dey and Joshi 2006). Lower *FI* values indicate greater constancy.

**Persistence**: Calculated as mean extinction frequency, EF = *E / T*, where *E* is the number of extinction events (metapopulation reaching zero individuals) across *T* generations. Lower EF indicates higher persistence.

**Genetic Stability (Effective Population Size)**: The *effective population size* (*N_e_*) was calculated as the harmonic mean of breeding metapopulation sizes: *N_e_ = T* / Σ(*1/N_t_*), where the sum is taken over *t* = 1 to *T*. Higher *N_e_* values reflect greater genetic stability and reduced drift. This metric is appropriate in our study because the unperturbed dynamics across the parameter ranges exhibit substantial fluctuations, and genetic drift is driven mainly by these repeated population bottlenecks. Even when control methods were applied, fluctuations were reduced but not eliminated, so *N_e_* continues to capture meaningful differences in population bottleneck severity. In inherently stable regimes, where control would not be required, *N_e_*-based comparisons are less relevant, and these lie outside the scope of this work.

**Synchrony**: Synchrony between subpopulations was quantified as the correlation coefficient at lag zero of first-differenced, log-transformed abundances of the two subpopulations (Bjørnstad et al. 1999; Dey and Joshi 2006). Synchrony values range from −1 to 1; positive values indicate synchronous fluctuations and thus greater risk of simultaneous subpopulation crashes.

### 2.4. Population control methods

We compared six rule-based population control methods: Constant Pinning (CP; McCallum 1992; Parthasarathy and Sinha 1995; Solé et al. 1999), Lower Limiter Control (LLC; Hilker and Westerho 2005), Adaptive Limiter Control (ALC; Sah et al. 2013), Upper Limiter Control (ULC; Hilker and Westerho 2005), Target-Oriented Control (TOC; Dattani et al. 2011), and Both Limiter Control (BLC; Tung et al. 2014). These methods represent the major strategies for stabilizing biological populations, including constant additions (CP), threshold-based restocking (LLC, ALC), threshold-based culling (ULC), proportional corrections toward a target (TOC), and dual-limit interventions (BLC). For each patch, each method modifies the post-dispersal population size (*b_t, i_*) according to its respective control rule to yield a post-control value (*a_t, i_*). Evaluating these methods within a unified metapopulation framework captures the main dimensions of intervention, including magnitude, direction (addition versus removal), and adaptiveness. These methods were previously analysed for isolated populations (Tung et al. 2014). Detailed mathematical formulations and parameter ranges are provided in Table 1.

**Table 1:** Mathematical formulation of the six control methods and their parameter range. Here, *b_t, i_* and *a_t, i_* are population sizes before and after external perturbations in patch *i* at time *t* respectively. *max*() and *min*() indicate maximum and minimum function respectively.

| Sl. No. | Control Method | Brief Description | Mathematical Expression | Control Parameters | Control Parameter Range | Step Size |
| --- | --- | --- | --- | --- | --- | --- |
| 1. | Constant Pinning (CP) | Add a constant number ($p$) of individuals at every time step | $a_{t,i} = b_{t,i} + p$ | Pin ($p$) | $[1, K - 1]$ | 1 |
| 2. | Lower Limiter Control (LLC) | Increase population size to the lower threshold $h$ whenever it falls below $h$ | $a_{t,i} = \max(b_{t,i}, h)$ | Lower limit ($h$) | $[1, K - 1]$ | 1 |
| 3. | Adaptive Limiter Control (ALC) | Similar to LLC, but with the lower threshold set as a fixed fraction ($c$) | $a_{t,i} = \max(b_{t,i}, c \times a_{t-1,i})$ | ALC intensity ($c$) | $[0.05, 0.95]$ | 0.05 |
| | | of the previous generation's breeding population size ($a_{t-1}$) | | | | |
| 4. | Upper Limiter Control (ULC) | Decrease population size to the upper threshold $H$ whenever it goes above $H$ | $a_{t,i} = \min(b_{t,i}, H)$ | Upper limit ($H$) | $[K+1, 3K]$ | 1 |
| 5. | Target Oriented Control (TOC) | Limit the population size within a predefined $c_d$ fraction of the target threshold $T$ (Here, the target is fixed at the carrying capacity, $K$) | $a_{t,i} = \max(0, b_{t,i} + c_d \times (T - b_{t,i}))$ | Target ($T$) | $K$ | Kept constant |
| | | | | TOC intensity ($c_d$) | $[0.05, 0.95]$ | 0.05 |
| 6. | Both Limiter Control (BLC) | A combination of LLC and ULC | $a_{t,i} = \max(h, \min(b_{t,i}, H))$ | Lower limit ($h$) | $[1, K - 1]$ | 1 |
| | | | | Upper limit ($H$) | $[K+1, 3K]$ | 1 |

### 2.5. Effort Magnitude (Cost)

Implementation cost was quantified as *effort magnitude (EM)*, defined as *EM =* Σ*|a_t_ − b_t_*| / (*N × T)*, where *b_t_* is metapopulation size before applying control, *a_t_* is metapopulation size after control, *N* is average metapopulation size (computed from the time series of *a_t_* values), *T* is the number of generations, and the sum is taken over *t* = 1 to *T*. This metric measures the average absolute net change in metapopulation abundance per generation relative to the realized metapopulation size maintained under that control method, thereby providing a dimensionless measure of net external intervention effort across regimes. We assumed equal unit cost for culling and restocking, consistent with prior theoretical frameworks (Tung et al. 2014, 2016a,b), so that observed differences in performance reflect the intrinsic effects of the control rules rather than cost asymmetries. In instances where one patch is augmented while the other is culled, EM captures the absolute net change in metapopulation size, assuming that individuals can be redistributed between patches. Alternatively, summing the absolute changes across individual patches would provide a measure of gross intervention effort.

### 2.6. The comparative framework and Composite Performance Score

For evaluating control methods, we used four combinations of intrinsic growth rate (*r*) and carrying capacity (*K*): 2.8 (low) and 4 (high) for *r*, and 60 (low) and 300 (high) for *K*. These values were selected to match those used in the single-population framework (Tung et al. 2014), enabling qualitative comparison between single-population and metapopulation results.

We examined three dispersal scenarios:

- Low symmetric dispersal: *m = m = 0.1*
- High symmetric dispersal: *m = m = 0.3*
- Asymmetric dispersal: *m = 0.1*, *m = 0.3*

Together, the four *r–K* combinations and three dispersal regimes produced 12 distinct ecological scenarios, capturing a wide range of demographic and spatial conditions under which the six control methods were compared.

Following the single-population study (Tung et al. 2014), we first identified, for each control method, the control parameter set that yielded a reduction in either fluctuation index (*FI*; constancy-targeted analysis) or mean extinction frequency (*EF*; persistence-targeted analysis) numerically closest to 50%. For each parameter combination, 70 replicate simulations were performed; no ties occurred under this procedure. For each goal, we compared methods using a Composite Performance Score (*CPS*) that integrates multiple management metrics into a single index. The composite performance score assigns equal weight to three components:

1. the non-target stability metric (*FI* or *EF*),
2. effective population size (*N_e_*), and
3. effort magnitude (*EM*, representing implementation cost).

These three components of the composite performance score operate on different numerical scales, which would otherwise bias comparisons. To avoid this, each component value was normalized by dividing it by the maximum value of that component among all six methods, separately for each of the 12 ecological scenarios. These normalized terms, referred to as component indices (*CI*), range from 0 to 1. The composite score was then computed as: *CPS*=*CI_FI/EF_*+*CI_EM_*+(1−*CI_Ne_*); where *CI_FI/EF_*represents the normalized value of either fluctuation index (for persistence-targeted analysis) or mean extinction frequency (for constancy-targeted analysis), CI_EM_ represents normalized effort, and *CI_Ne_* represents normalized effective population size. Lower composite performance score values indicate better overall performance. Because a higher *N_e_* is beneficial, its contribution is inverted as (1−*CI_Ne_*) in the equation.

In low-growth regimes (*r*=2.8), mean extinction frequency was nearly zero even without intervention, making persistence an irrelevant management target. This reflects the fact that low-r Ricker dynamics exhibit weaker overcompensation and thus rarely generate the severe crashes required to trigger lattice-effect or demographic-stochasticity–driven extinctions.

Therefore, *CI_EF_*was excluded from composite performance score calculations for constancy-targeted analyses in these regimes to avoid overemphasizing non-existent extinction risk.

This composite framework provides a balanced and standardized means of comparing rule-based control strategies across diverse ecological contexts, ensuring that trade-offs among stability, persistence, genetic health, and management cost are evaluated on equal footing.

### 2.7. Robustness to Census Error

All the above control methods except Constant Pinning rely on accurate population size estimates each generation. In natural settings, population censuses are inherently noisy due to detection errors, observer bias, and environmental variability affecting detectability, among other factors. To evaluate the robustness of control methods under such census uncertainty, we extended our analysis to compare performance scores in the presence of observation noise.

We modelled three classes of census error applied independently to the true post-dispersal population size of each patch (*b_t, i_*):

1. White noise (unbiased error): *N_obs, i_ = b_t, i_ + U(−*ε *× b_t, i_,* ε *× b_t, i_)*;
2. Positive bias (overestimation): *N_obs, i_ = b_t, i_ + U(0,* ε *× b_t, i_)*;
3. Negative bias (underestimation): *N_obs, i_ = b_t, i_ + U(−*ε *× b_t, i_, 0)*,

where *U* denotes a uniform distribution and ε is the error level. Error levels (ε) ranged from 0 to 0.5 in increments of 0.05, representing 0–50% deviation from the true population size. In all simulations, control decisions were made based on the observed population size (*N_obs, i_*), while demographic updates followed from the true size (*b_t, i_*), reflecting realistic management conditions. The control parameter values themselves were not re-estimated for each error level; for each method and ecological scenario, the same parameter set identified under the corresponding zero-error analysis was used across all census error levels.

We first computed composite performance scores (CPS; as per section 2.6) separately for each census error level, with component normalization performed independently for each ecological scenario at that error level, and then averaged CPS over 20 independent repetitions to account for stochastic variability. Consequently, CPS values at different error levels are interpretable in terms of the relative performance of methods within that error level, rather than absolute performance on a common normalized scale. We therefore assessed robustness to census error by examining how mean CPS trajectories and the rank ordering of methods were preserved across error levels.

### 2.8. Simulation Protocol and Replication

At the start of each simulation, all replicates in every treatment were initialized with 20 individuals per subpopulation. When the total metapopulation size reached zero, both subpopulations were reset to eight individuals, to allow continued simulation and evaluation of control methods over extended time series. This reset protocol follows the single-population study (Tung et al. 2014) and is standard in theoretical metapopulation studies where multiple stability metrics must be estimated across replicates (Dey and Joshi 2006; Sah et al. 2013; Tung et al. 2019). Without resets, replicates that reach extinction would terminate permanently, precluding estimation of constancy, effective population size, and control method performance in extinction-prone regimes. Sensitivity analyses across a range of initial and reset values (Supplementary Text ST1) confirmed that these choices did not qualitatively affect the results (Supplementary Figures S1-2). Because the same reset rule applies identically to all treatments including the uncontrolled baseline, it maintains simulation continuity without biasing comparisons among control methods.

For baseline (uncontrolled) dynamics, each parameter combination was simulated for 50 independent replicates to account for stochastic variation, and mean stability metrics were recorded. For simulations involving control methods, 70 independent simulation replicates were used to estimate mean stability components for each method. This entire composite performance score (*CPS*) calculation procedure was then repeated 20 times, and *CPS* values were averaged across repetitions. Results are reported as means ± standard error of the mean (SEM).

For each generation, the following sequence of events was implemented in both subpopulations:

1. Introduce environmental stochasticity to *r* and *K*.
2. Apply Ricker dynamics to update population size.
3. Apply demographic stochasticity (50% extinction probability when *N* < 4).
4. Implement dispersal between patches, producing the post-dispersal population sizes (*b_, i_*).
5. Add census noise to *b _, i_* to obtain *N_obs, i_* (only for robustness analyses); otherwise, *N_obs, i_* = *b_, i_*.
6. Apply the designated control method using observed population size (*N_obs, i_*) to determine the control action and yield *a_, i_*, the post-control population size.
7. Record *b _, i_* (or *N _bs, i_* for robustness analyses) and *a* to compute individuals added or removed, and update all stability metrics.

This workflow ensures that control decisions are based on post-dispersal abundances, aligning with realistic management timing where census and intervention occur after dispersal events.

### 2.9. Computational Implementation and Acronyms

These simulations were implemented using Python 3.8.16 (https://www.python.org/) and graphs were plotted using its *matplotlib* module. All scripts used in this work are available in the supplementary online materials. All Python scripts required to reproduce the simulations and analyses are included in the supplementary file *Scripts.rar*. For clarity, all acronyms used for stability metrics and performance measures are summarized in Table 2.

**Table 2.**
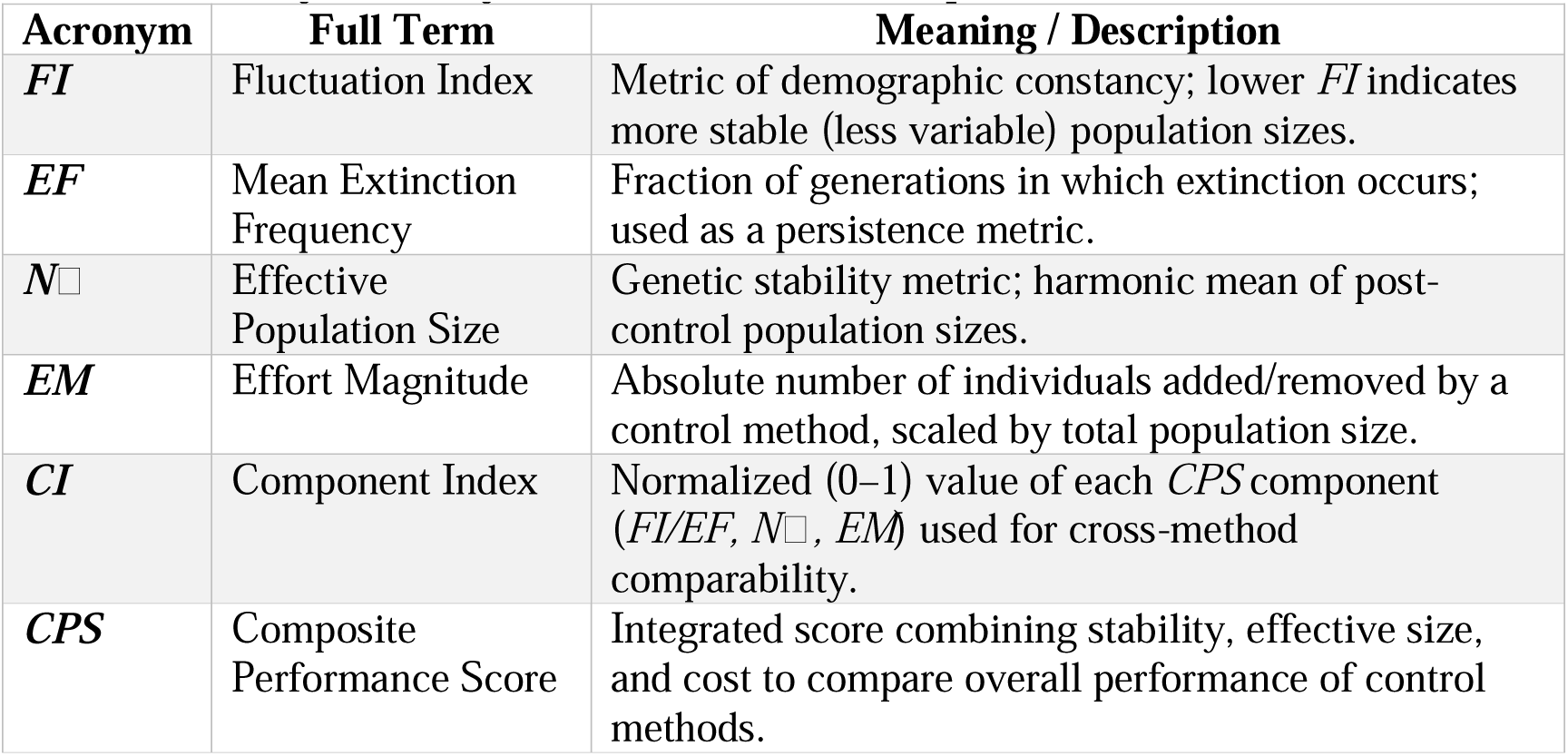
Glossary of acronyms used in the manuscript.

## 3. Results and Discussion

### 3.1. Dynamics of the Unperturbed Regimes

We first characterized metapopulation dynamics without control interventions across a broad grid of intrinsic growth rates (*r* = 2.7–4.5) and carrying capacities (*K* = 20–350) under low and high symmetric dispersal (*m = m* = 0.1, 0.3) and asymmetric dispersal. We quantified demographic constancy (fluctuation index, *FI*), persistence (mean extinction frequency, *EF*), and genetic stability (effective population size, *N*) of the metapopulation, thereby establishing a baseline for subsequent comparisons.

Regimes with high *r* and low *K* were consistently the most unstable, exhibiting large-amplitude boom–bust cycles, frequent population bottlenecks, and elevated extinction risk (Figure 2), which are classic signatures of overcompensatory dynamics in discrete-time systems (May and Oster 1976; Grimm and Wissel 1997; Tung et al. 2014). Repeated crashes depressed *N* because the harmonic mean is dominated by low counts, indicating strong drift and potential inbreeding despite occasional high abundances (Allendorf et al. 2012). These qualitative patterns held under both symmetric and asymmetric dispersal.

**Figure 2.**
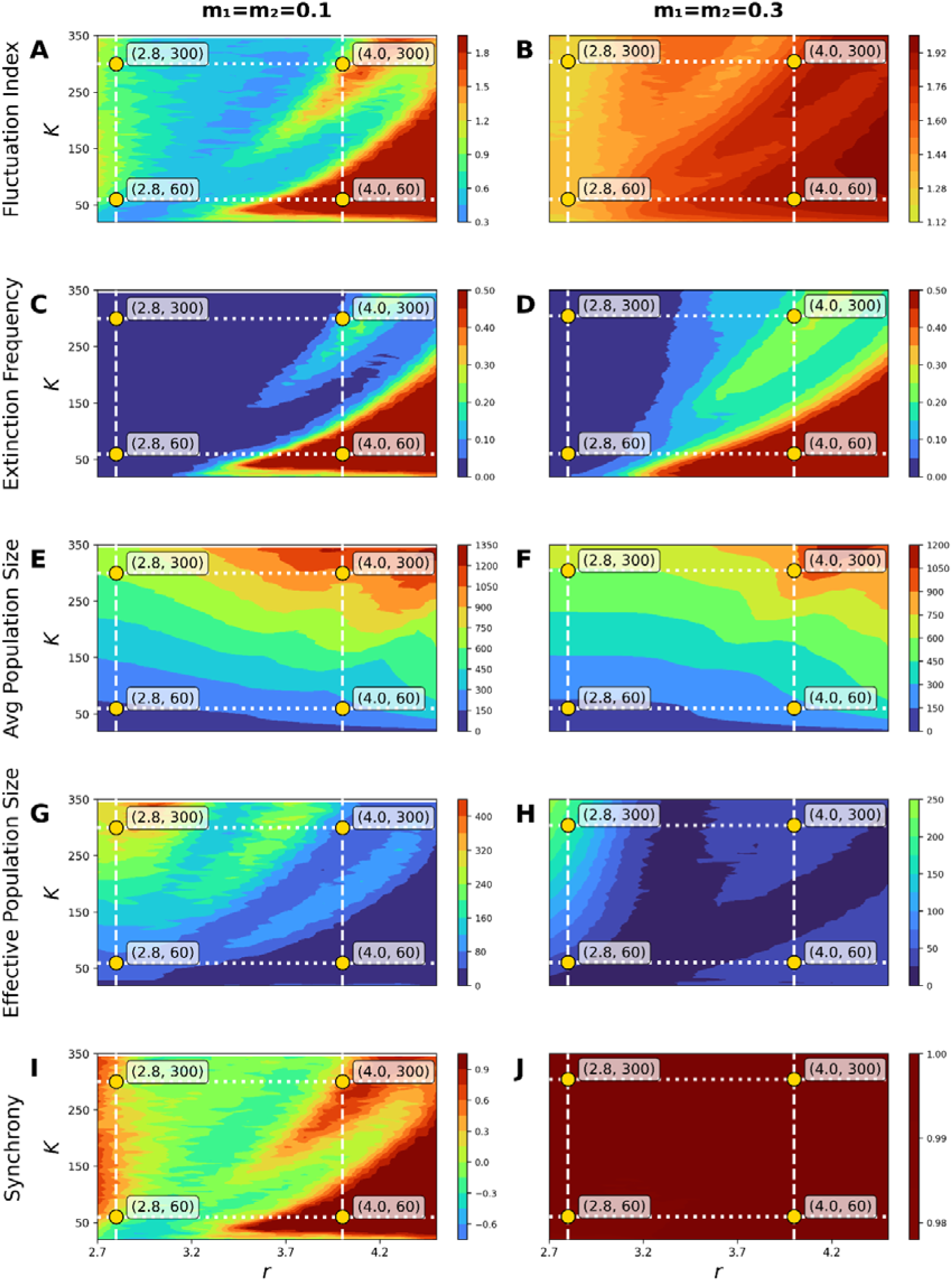
Unperturbed Metapopulation Dynamics. Heatmaps show how key metapopulation metrics vary across the *r–K* parameter space for two symmetric dispersal rates (left: m = 0.1 (low); right: m = 0.3 (high)). Warmer colours indicate higher values. (A,B) fluctuation index, (C,D) extinction frequency, (E,F) average metapopulation size, (G,H) effective population size, and (I,J) spatial synchrony. White dashed boxes denote the focal *r–K* values (2.8, 4.0; 60, 300), and yellow circles mark four focal parameter combinations analysed for comparing the performance of control methods.

Regimes with high *r* and high *K* also fluctuated strongly but showed lower *EF* than their low-*K* counterparts, as larger carrying capacities buffered complete collapse (Tung et al. 2014). Nevertheless, substantial oscillations still limited *N* for the same harmonic-mean reason (Allendorf et al. 2012). Notably, high-*K* regimes often showed high mean population sizes but relatively low *N_e_*, highlighting that large carrying capacity does not simply inflate *N_e_* proportionally, because deep crashes strongly depress the harmonic mean. Dispersal level modulated constancy via synchrony: low symmetric dispersal (*m = m* = 0.1) maintained out-of-phase subpopulation cycles and reduced metapopulation FI, whereas high symmetric dispersal (*m = m* = 0.3) synchronized dynamics and amplified metapopulation-level variation—an effect predicted by theory and observed empirically (Gyllenberg et al. 1993; Doebeli 1995; Amarasekare 1998; Kendall and Fox 1998; Mueller and Joshi 2000; Ylikarjula et al. 2000; Briggs and Hoopes 2004; Lecomte et al. 2004; Dey and Joshi 2006; Abbott 2011; Sah et al. 2013; Dey et al. 2014).

By contrast, low**-***r* regimes exhibited greater demographic stability: *FI* was low and *EF* was near zero across dispersal scenarios, with especially strong constancy under low symmetric dispersal where asynchrony reduced variance at the metapopulation scale (Dey and Joshi 2006). Genetic outcomes diverged with resource level: low**-***r*, low**-***K* regimes were demographically stable yet maintained small *N* due to persistently low census sizes, whereas low**-***r*, high**-***K* regimes combined high persistence and constancy with the largest *N* values. Across the landscape, high extinction risk was typically associated with low genetic diversity proxies (sensu DiLeo et al. 2024), although the converse need not hold.

Under asymmetric dispersal (Supplementary Figure S3), metapopulation stability generally fell between the two symmetric cases. Synchrony remained low only when dispersal rates were low in both directions, whereas high migration from either patch rapidly synchronized population dynamics. This pattern shows that even partial directional connectivity can strongly couple subpopulations, eroding the variance-buffering advantage of spatial asynchrony, which is a hallmark of nonlinear coupling dynamics (Doebeli 1995; Ylikarjula et al. 2000; Dey and Joshi 2013). In the low**-***r*, low**-***K* regime, however, extreme one-way dispersal from either patch slightly increased mean metapopulation size, likely because strong unidirectional flow acted as a rescue mechanism, sustaining low-productivity patches through continuous immigration without inducing extinction due to demographic stochasticity (EDS; Tung et al. 2014). Such demographic rescue effects disappeared under high-*r* conditions, where stronger coupling dominated the dynamics.

To balance ecological generality with analytical tractability, we evaluated control methods across four representative regimes (*r* = 2.8, 4.0; *K* = 60, 300), capturing contrasting demographic–genetic conditions while maintaining comparability with the single-population framework (Tung et al. 2014). Each regime was examined under three dispersal scenarios: low symmetric (m_1_ = m_2_ = 0.1), high symmetric (m_1_ = m_2_ = 0.3), and asymmetric (m_1_ = 0.1, m_2_ = 0.3), yielding 12 ecologically distinct scenarios.

### 3.2 Comparing constancy-targeted control methods

Our findings indicate that the relative performance of control methods in metapopulations is context-dependent, with no single method proving universally optimal across all regimes (Figure 3). This echoes previous observations in spatially unstructured populations (Tung et al. 2014). Interestingly, our comparative analysis identifies a common trend for constancy-targeted control in regimes with low intrinsic growth rate (i.e. low-*r* regimes). In these regimes, lower limiter control (LLC), adaptive limiter control (ALC) and both limiter control (BLC) demonstrated similar composite scores and consistently outperformed other methods (typically ranking 1–3 across these regimes; Figure 3A-D and Figure S4A-B). These are closely followed by target-oriented control (TOC) and upper limiter control (ULC), in the order of their performance. Constant pinning (CP) method performed consistently worst in all scenarios. The reason behind this trend became clear when we checked the individual components (Figure S6-S8). In low-*r* regimes, effective population size and effort magnitude constituted the composite score. LLC, ALC, and BLC achieved effective population sizes that were comparable and only lower than those induced by the CP method. Although CP had the potential to maximize effective population size across the regimes, its high implementation costs, as evidenced by substantial effort magnitude, made it the least favourable option. In contrast, LLC, ALC, and BLC struck a balance, attaining moderate effective population sizes with reasonable level of implementation efforts to become the best performing methods in these regimes.

**Figure 3.**
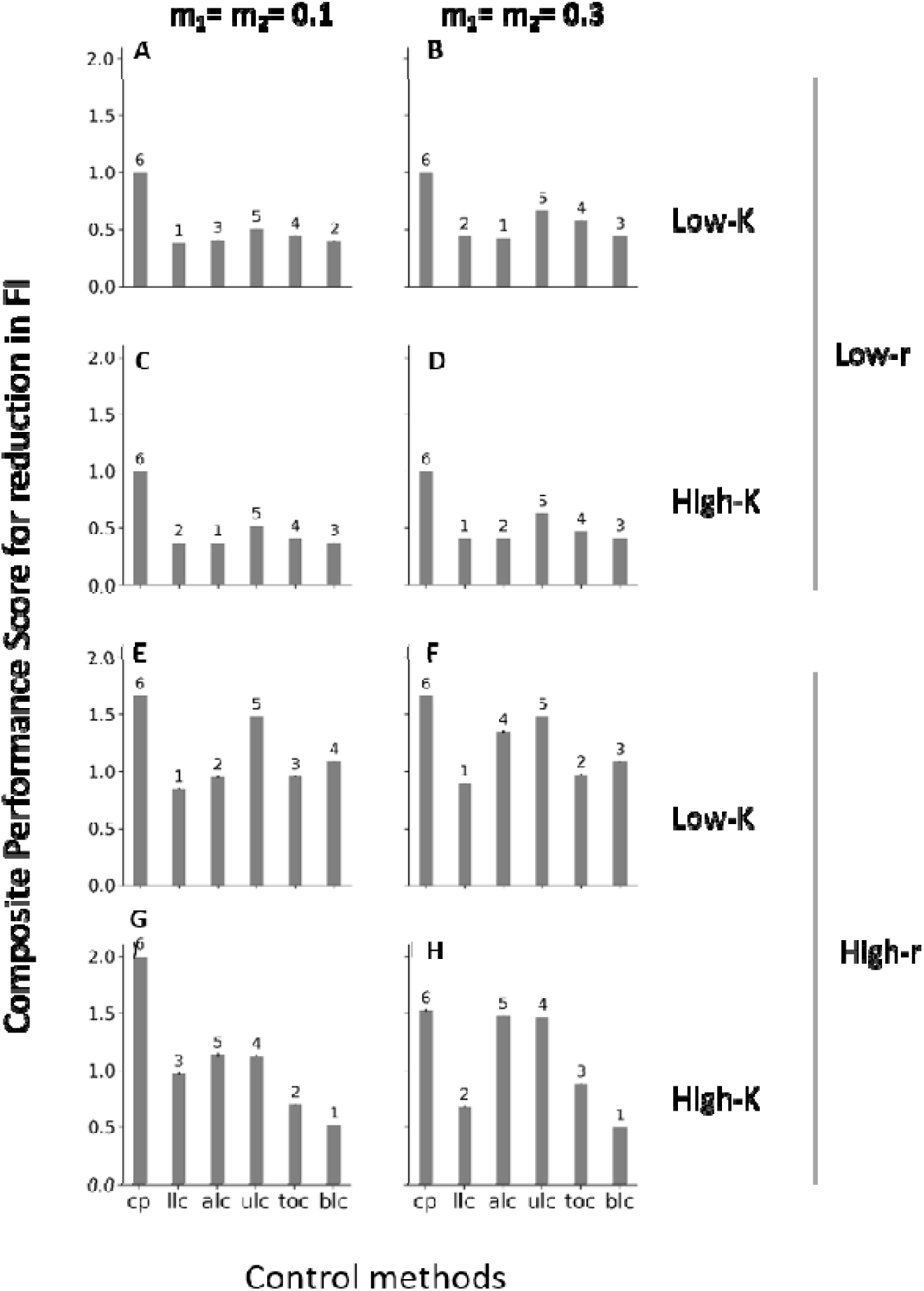
Composite Performance Scores for 50% Reduction in Fluctuation Index. Presented here are the composite performance scores for six population control methods evaluated across eight distinct regimes with low (A, C, E, G) and high (B, D, F, H) symmetric dispersal, with the target of achieving a 50% decrease in Fluctuation Index to promote constancy stability. See Supplementary Figure S4 for corresponding graphs with asymmetric dispersal rates. Atop each bar, the numerical annotations represent the performance ranking of the method, with a lower composite score denoting a superior rank. Standard error is depicted as the error bars; however, in most instances, the error bars are too small to be visible.

In high-*r*, low-*K* regimes, LLC stood out as the superior method (rank 1 across these regimes; Figure 3E-F and Figure S4C). It paralleled ALC in enhancing effective population size. However, LLC surpassed ALC in reducing implementation effort and mean extinction frequency, establishing itself as the most favourable method in these scenarios. In constancy-targeted analysis, while ULC and BLC proved more favourable in mitigating extinction risks, their higher implementation costs and lower effective population sizes negatively impacted their overall rankings according to the composite scores. On the other hand, in high-*r*, high-*K* regimes, BLC performed favourably (rank 1 in these regimes), successfully eliminating extinction risks while achieving an optimal balance between effort and effective population size (Figure 3G-3H and Figure S4D).

### 3.3 Comparing persistence-targeted control methods

In our subsequent analysis, we focused on the performance of control methods that were designed to reduce the mean extinction frequency to 50% of the corresponding unperturbed scenario. Since low-r regimes rarely experience extinction, this analysis was restricted to high-r regimes. Here, the composite performance score was calculated from the fluctuation index, effective population size, and the magnitude of implementation effort required for reducing the mean extinction frequency by 50% (following Tung et al. 2014).

We found that ALC was the most favourable method across all high-*r* regimes under the persistence-targeted analysis (rank 1 throughout; see Figure 4 and Figure S5), closely followed by LLC, BLC, and CP. In contrast, ULC and TOC consistently ranked less favourably. This suggests that methods designed to prevent population size from dropping below a certain threshold – whether constant (as in CP, LLC and BLC) or variable (as in ALC) – tended to rank more favourably overall under the persistence-targeted framework than methods based primarily on culling overabundant populations, such as TOC and ULC.

**Figure 4.**
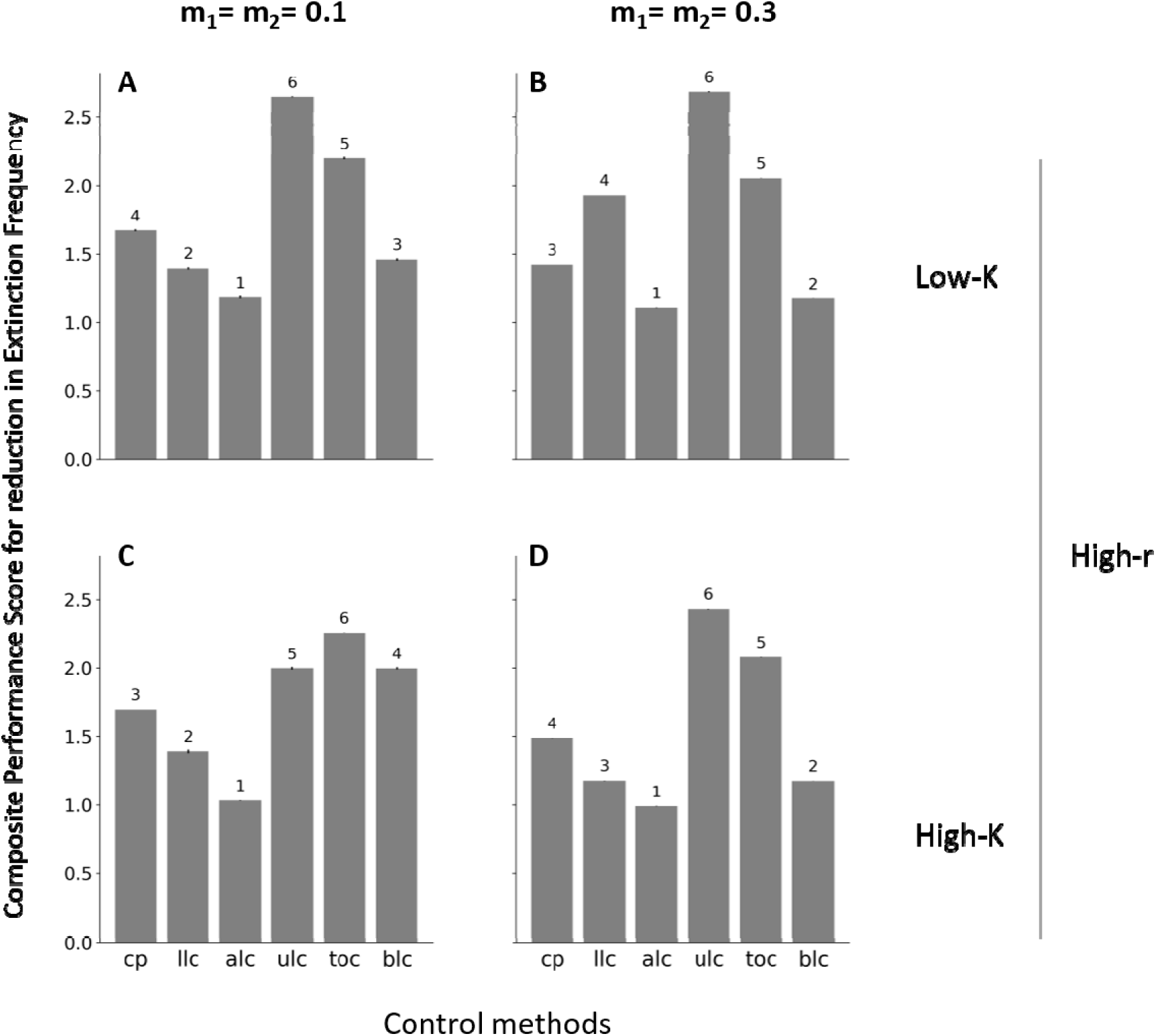
Composite Performance Scores for 50% Reduction in Extinction Probability. Presented here are the composite performance scores for six population control methods evaluated across four regimes, with high intrinsic growth rate (i.e. High-*r* regimes) with symmetric low (A, C) and high (B, D) dispersal rate, while achieving a 50% decrease in Extinction Probability to promote persistence stability. See Supplementary Figure S5 for corresponding graphs with asymmetric dispersal rates. Atop each bar, the numerical annotations represent the performance ranking of the method, with a lower composite score denoting a superior rank. Standard error is depicted as the error bars; however, for most instances, the error bars are too small to be visible.

Interestingly, although ALC ranked best under persistence-targeted analyses across all regimes, examination of the CPS components revealed that it achieved this favourable overall performance through different routes (Figure S9-S11). In case of the most unstable high-*r*, low-*K* regimes, when migration rate between the subpopulations was low (i.e. for high-*r*, low-*K* regime with symmetric low dispersal), CP induces the maximum effective population size followed by LLC, ALC and BLC. But CP does that at the cost of a large effort magnitude leading to its poor performance rank based on composite score. Among LLC, ALC and BLC, ALC performed the best with respect to curbing population size fluctuation and reducing effort magnitude, leading to low composite scores that identified it as the best-performing method overall under the persistence-targeted framework in this regime. In scenarios with high migration rate between the subpopulations i.e. in high-*r*, low-*K* regime with symmetric high dispersal, LLC had the smallest effort magnitude but was associated with very low effective population size, compromising its composite performance score.

Other trends were similar, and overall ALC ranked most favourably under the persistence-targeted framework. When migration between the subpopulations was asymmetric i.e. in high-*r*, low-*K* regime with asymmetric dispersal, the effort magnitude associated with ALC under the persistence-targeted framework was actually the maximum, but as this method was excellent in simultaneously improving constancy stability and genetic stability, it therefore ranked most favourably overall, with the lowest composite performance score. When carrying capacity was high, ALC combined low population-size fluctuation and high effective population size with only moderate effort magnitude, yielding the most favourable overall composite score. While LLC and BLC also ranked favourably, they fell short compared to ALC in one or more components of the composite performance score.

### 3.4 Summary of constancy- and persistence-targeted comparisons

To provide a consolidated overview of method performance across both constancy- and persistence-targeted analyses, we compiled a summary table aggregating method ranking across all evaluated regimes (Table S1). This synthesis indicates that Lower Limiter Control (LLC) most consistently ranked favourably under the constancy-targeted framework, combining high effective population size, strong persistence, and low effort. Under the persistence-targeted framework, Adaptive Limiter Control (ALC) ranked most favourably overall across all evaluated regimes (Table S1), maintaining advantageous combinations of effective population size, extinction risk, and cost. These results summarize comparative method performance across ecological contexts in spatially structured populations and highlight the approaches most consistently associated with favourable outcomes, thereby informing the choice of control strategies in metapopulations.

### 3.5 Comparing robustness of performance rankings of the control methods against census noise

In constancy-targeted analyses, the effects of census noise were more clearly structured by carrying capacity than by intrinsic growth rate (Figure 5; Supplementary Figure S12). In low-K regimes, the ranking of control methods changed little across census error levels and was broadly similar for both low-r and high-r dynamics (Figure 5A, B, E, F; Supplementary Figure S12A, C). CP consistently remained the worst-ranked method, ULC the second worst, TOC occupied an intermediate position, and LLC, ALC, and BLC consistently retained the least composite performance scores (CPS) and therefore, most favourable comparative ranks (Figure 5A, B, E, F). Thus, in low-K regimes, the main effect of census noise was not to reorder method choice, but to preserve a broadly stable separation between the poorer-performing methods (CP and ULC) and the better-performing methods (LLC, ALC, and BLC).

**Figure 5.**
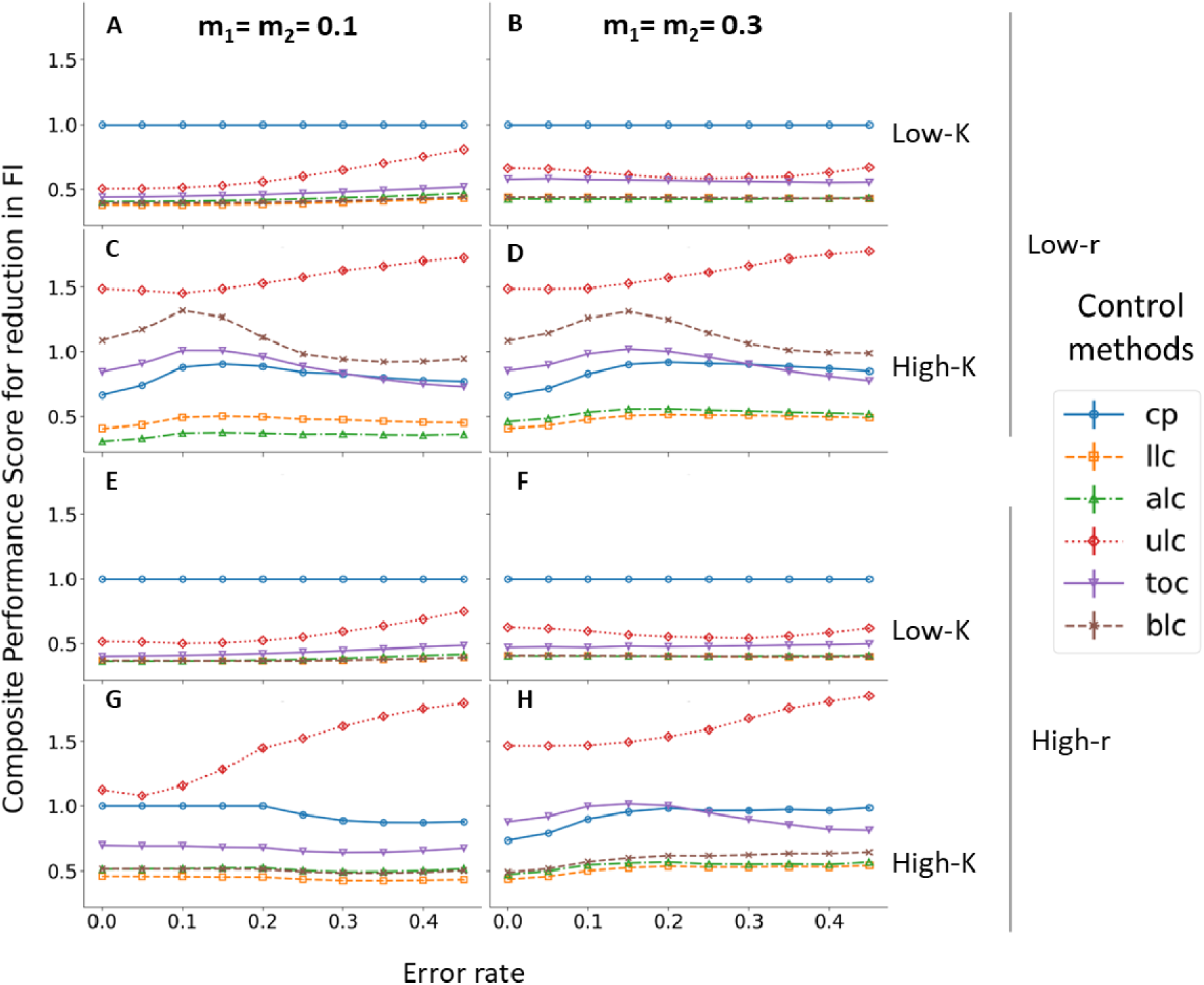
Change in average composite scores for reducing Fluctuation Index (*FI*) with change in intensity of white noise for regimes with symmetric low (=0.1) and high (=0.3) dispersal. Corresponding graph for scenarios with asymmetric dispersal rates can be found in supplementary Figure S12. Note for computing composite performance score, only effort magnitude (*EM*) and effective population size (*N_e_*) were used to calculate the composite score for low-r regimes, since average extinction frequencies (*EF*) were negligible even without any perturbations. For high-*r* regimes, however, extinction frequency was included in calculating the composite score. The vertical lines on the line plots denote standard error around average composite scores across the replicate runs.

In high-*K* regimes, a different pattern emerged. Here, ULC consistently occupied the poorest comparative rank across census error levels, while LLC and ALC remained among the best-ranked methods (Figure 5C, D, G, H; Supplementary Figure S12B, D). The remaining methods were more regime dependent. Under low-*r*, high-*K* conditions, BLC showed a pronounced non-monotonic CPS trajectory and was less favourably ranked than CP and TOC over the error range (Figure 5C, D). Under high-*r*, high-*K* conditions, however, BLC occupied a more favourable comparative position, while CP and TOC showed somewhat greater shifts in relative rank with increasing noise (Figure 5G, H). Taken together, these results indicate that, across high-*K* regimes, the most consistent pattern under census noise was the preservation of favourable relative ranks for LLC and ALC and unfavourable relative ranks for ULC, whereas CP, TOC, and BLC accounted for most of the remaining regime-specific variation in ranking stability.

In persistence-targeted analyses, under symmetric migration, ALC consistently retained the most favourable comparative rank across census error levels and dispersal scenarios (Figure 6). In high-*r*, low-*K* regimes, the ranking of methods changed little overall, although some reordering occurred among the poorer-performing methods; ULC and TOC generally occupied unfavourable ranks, with ULC showing the largest shifts in relative position (Figure 6A, B). In high-*r*, high-*K* regimes, method rankings were more consistently preserved across error levels, with ALC remaining best ranked throughout, LLC or BLC typically occupying the next most favourable positions, CP remaining intermediate, and ULC and TOC generally ranking poorly (Figure 6C, D). Under asymmetric migration as well, ALC remained the best-ranked method, closely followed by LLC (Supplementary Figure S13). Taken together, these patterns indicate that ALC showed the most consistently favourable and stable relative performance for persistence-targeted analysis under census noise.

**Figure 6.**
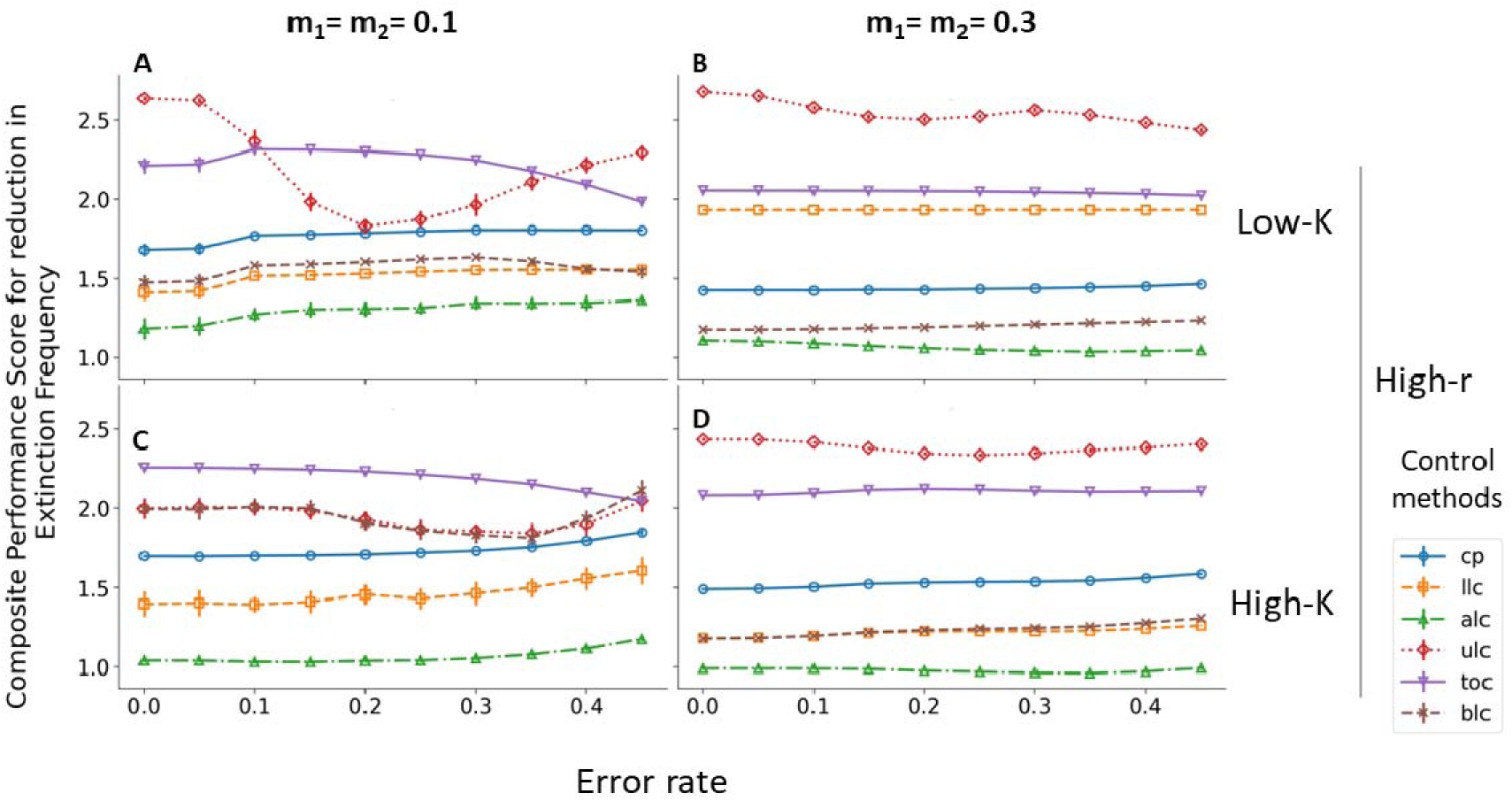
Change in average composite scores for reducing Extinction Frequency (*EF*) with change in intensity of white noise for regimes with symmetric low (=0.1) and high (=0.3) dispersal. Corresponding graph for scenarios with asymmetric dispersal rates can be found in supplementary Figure S14. The label on the y axis indicates that Fluctuation index (*FI*), Effort magnitude (*EM*), and Effective population size (*N_e_*) were used to calculate the composite score. The vertical lines on the line plots denote standard error around average composite scores across the replicate runs.

Overall, ALC retained favourable comparative ranks across a range of intensities of positively biased (i.e. overestimation; Supplementary Figures S14–S15) and negatively biased (i.e. underestimation; Supplementary Figures S16–S17) census noise. By contrast, ULC showed the greatest variation in composite score across noise intensities. Differences in the stability of relative performance across control methods appear to arise from how each method uses census information (Supplementary Figures S14–S17). Lower-threshold methods (LLC, ALC, and BLC) intervene only when populations fall below a critical level; when populations remain well above this threshold, small census errors are less likely to alter intervention decisions. ALC is additionally buffered because its threshold is defined relative to the previous census, so correlated noise across successive measurements tends to preserve the relative comparison that triggers intervention. In contrast, ULC depends on accurate detection of upper-threshold crossings, making its relative performance particularly sensitive to census error. Under positively biased noise, ULC showed a distinct U-shaped response, with more favourable relative composite scores at intermediate noise intensities (Supplementary Figures S14–S15). A plausible explanation is that moderate overestimation causes observed population sizes to cross the fixed upper threshold at slightly lower true abundances, thereby inducing earlier culling and reducing extreme peaks in high-*r* regimes. At high overestimation intensities, however, census estimates became too unreliable for coherent control, and ULC’s relative position again became less favourable. This pattern was not observed under negatively biased noise (Supplementary Figures S16–S17), where the operative thresholds were raised and control weakened without producing a comparable intermediate optimum. Taken together, these patterns suggest that the distinct response of ULC under positively biased noise reflects a method-specific interaction between census bias and threshold-based intervention timing. CP did not show an analogous mechanistic response: although its control rule is independent of census information and its control parameter was held constant across error levels, its CPS could still vary because CPS components were normalized separately within each error level. Consequently, changes in CPS for CP reflect shifts in its relative standing among methods under a given error level, rather than a direct effect of census noise on the CP rule itself.

### 3.6 Metapopulation vs. Single-Population Control Outcomes: A Comparative Synthesis

Our comparative analysis shows that, contrary to our initial predictions, dispersal does not invert the hierarchy of control methods established in the single-population study (Tung et al. 2014); instead, augmentation-based strategies remain the most favourable in spatially structured metapopulations. Lower Limiter Control (LLC) consistently provides the strongest improvements in constancy stability across most dispersal scenarios, suggesting that the fundamental advantage of trough-elevating interventions under high-*r* dynamics persists whether populations are isolated or coupled. Adaptive Limiter Control (ALC) emerges as the most favourable method for enhancing persistence across all high-*r* regimes, irrespective of dispersal intensity or symmetry, with its proportional threshold allowing it to perform reliably across spatial contexts and to maintain a consistently favourable relative ranking across census noise levels. Both Limiter Control (BLC), which combines restocking and culling, becomes increasingly competitive at high carrying capacities, but does not surpass ALC in persistence outcomes. In contrast, culling-dominated approaches such as Upper Limiter Control (ULC) and Target-Oriented Control (TOC) do not gain an advantage under high dispersal. Instead of mitigating dispersal-induced overshoots as initially anticipated, these methods perform poorly because they do not protect against synchrony-driven crashes and depressed harmonic-mean population sizes in metapopulation settings. Constant Pinning (CP) maximizes *N_e_* in isolated populations and metapopulations alike, but its consistently high implementation effort reduces its overall utility across all dispersal conditions. Collectively, these results indicate that, under the parameter ranges explored, the control strategies identified as most favourable in isolated populations remain broadly well suited to controlling metapopulations. At the same time, the metapopulation context clarifies why this ranking is preserved: overcompensatory growth and demographic stochasticity, rather than overshoot-driven collapses alone, appear to be the dominant sources of instability in these spatially structured systems. Consequently, interventions that prevent populations from approaching low densities outperform methods aimed primarily at limiting high densities. These findings reinforce theoretical expectations that dispersal acts as a double-edged sword, simultaneously damping local variability while amplifying spatial synchrony (Gyllenberg et al. 1993; Hanski 1998; Dey and Joshi 2006), and highlight the importance of evaluating population-stabilization strategies within spatially realistic metapopulation frameworks rather than extrapolating uncritically from isolated populations.

### 3.7 Caveats

This study presents a general framework for identifying population control methods capable of stabilizing metapopulations whose subpopulation dynamics follow the two-parameter Ricker model. Our approach involves specifying the metapopulation structure (here, two subpopulations connected by bidirectional migration), defining local dynamics (Ricker growth followed by dispersal), applying control methods to achieve a target stability criterion (e.g., 50% reduction in *FI* or *EF*), and comparing the resulting controlled systems using a composite performance score (*CPS*) based on additional stability metrics, implementation effort (EM), and effective population size (*N_e_*). The *CPS* therefore serves as a decision-support tool for method selection under different parameter regimes, both with and without census error.

Several aspects of our design define the scope within which these findings apply. First, our analysis is restricted to a two-patch, spatially homogeneous system. Increasing the number of patches or introducing spatial heterogeneity in growth parameters could alter synchrony, rescue effects, and the relative performance of control methods, especially those relying on fixed versus adaptive thresholds. Second, we evaluated methods using four representative *r– K* combinations, chosen to capture the contrasting dynamical regimes revealed by preliminary parameter sweeps and to retain direct comparability with single-population study (Tung et al. 2014). Extending the analysis across a broader *r–K* landscape may uncover additional context dependence. Third, the dominant mechanism of extinction in our model arises from intrinsic fluctuations that push populations below the extinction threshold, amplified by demographic stochasticity at low numbers (n ≤ 4). In many natural systems, however, extinctions may instead be driven by trait–environment mismatches, habitat degradation, or climate-induced variability in *r* and *K*. Consequently, our results apply specifically to internally driven extinctions and may not generalize to populations threatened primarily by external environmental forcing. Fourth, following extinction events, populations were reset to fixed values to preserve simulation continuity; although sensitivity analyses indicate limited influence within our parameter ranges, different reset rules could interact with specific control mechanisms and merit exploration in future work.

Our framework itself is highly flexible. Alternative growth models (e.g., Beverton and Holt 1993) can be incorporated by substituting the appropriate subpopulation dynamics. The definition of the CPS can likewise be adapted to management priorities: we did not include census effort explicitly, yet this cost varies among methods (no census for CP; minimal for LLC/ALC; threshold-dependent monitoring for ULC, TOC, and BLC). We assigned equal weights to stability, EM, and N_e_, but real-world conservation objectives may prioritize genetic diversity, persistence, or minimal intervention cost. Finally, the framework can be extended to larger or more complex metapopulation networks by modifying the dispersal structure and associated dynamical equations.

## 4. Conclusion

In conclusion, our comparative analysis of six methods for stabilizing metapopulation dynamics, along with their behaviour under census noise, reveals how demographic regime, carrying capacity, and migration structure jointly shape control outcomes. No single method emerged as universally superior across all ecological scenarios. Under the constancy-targeted framework, Lower Limiter Control (LLC), Adaptive Limiter Control (ALC), and Both Limiter Control (BLC) were the most favourable options in low-r regimes. In high-*r*, low-*K* regimes, LLC provided the best overall balance under the constancy-targeted framework, combining target achievement with relatively high effective population size and low implementation cost. In high-*r*, high-*K* regimes, BLC emerged as the best-performing method overall under the same framework. When persistence stability was the target, ALC consistently emerged as the strongest performer across ecological regimes, achieving the desired reduction in extinction frequency while maintaining high *N* and low effort, although the basis of this advantage varied with migration scenario.

Our findings emphasize that the choice of stabilization method must depend on which aspect of stability is being targeted. They further show that, under imperfect census data, the relative ranking of methods was often broadly preserved across noise levels, with LLC, ALC, and, in several constancy-targeted regimes, BLC frequently retaining favourable comparative positions. Thus, while census noise can modify comparative scores, it often does not overturn the overall choice of preferred method. Notably, in the persistence-targeted analyses, ALC consistently retained the most favourable comparative rank across census error levels, irrespective of intrinsic growth rate, carrying capacity, or migration scenarios. More broadly, this study advances our understanding of controlled metapopulation dynamics and provides a general framework for evaluating population stabilization strategies in spatially structured systems. It also highlights the need for a nuanced, context-dependent approach when selecting control methods for ecological management and conservation.

## Acknowledgement

Akanksha Singh thanks Department of Science and Technology, Government of India, for financial support through a KVPY fellowship. Sudipta Tung acknowledges the support of DBT/Wellcome Trust India Alliance Early Career Fellowship (#IA/E/18/1/504347) and Ashoka University.

## Authors’ contributions

Akanksha Singh and Sudipta Tung formulated the study. Akanksha Singh and Sudipta Tung carried out the simulations. Sudipta Tung and Akanksha Singh wrote the manuscript.

