## Supplementary for "Comparison of six methods for stabilizing metapopulation dynamics and the robustness of performance rankings to census noise"

for

**ST1. Sensitivity to initial conditions and reset values**

To evaluate whether our choice of initial population size (N₀ = (20, 20)) and post-extinction reset value (8, 8) might constrain simulations to particular transient dynamics, we conducted systematic sensitivity analyses. For initial population sizes, we simulated baseline metapopulation dynamics across N₀ values ranging from 0 to 350 (step size 5) for patch 1 while holding patch 2 at N₀ = 20, thereby creating strongly asymmetric initial conditions. For reset values, we varied the reset value for patch 1 from 0 to 15 while holding patch 2 constant at 8. In both analyses, we computed all four stability indices (fluctuation index, mean extinction frequency, effective population size, and synchrony) and average population size across the full range of r values (2.7–4.5) for both low (= 60) and high (= 300) K values.

Our sensitivity analyses demonstrate that stability indices are robust to initial and reset conditions (Supplementary Figures S1, S2). When we varied initial population size in patch 1 across a wide range (*N₀* = 0–350) while holding patch 2 constant, all five indices changed primarily with intrinsic growth rate (*r*) and were largely insensitive to *N₀*. This insensitivity arises because generation-wise stochastic variation in *r* and *K* causes trajectories to diverge rapidly, preventing simulations from remaining confined to particular transient paths despite identical starting conditions. Notably, synchrony remained consistently high across all *N₀* values, confirming that the synchronous dynamics we report are not artifacts of symmetric initial populations. For post-extinction reset values, sensitivity emerged only under a narrow set of conditions, specifically, when reset ≤ 3 and *r* > 3.7, where populations recolonizing from near-zero face elevated extinction risk before reaching stable dynamics. Our default reset value of 8 lies well outside this sensitive region.

**Table S1. Rank sums of composite scores of the six control methods for constancy- and persistence-targeted analysis.** Lower rank sum indicates better overall performance.

| **Analysis type** | **CP** | **LLC** | **ALC** | **ULC** | **TOC** | **BLC** |
| --- | --- | --- | --- | --- | --- | --- |
| Constancy-targeted analysis (across all 12 regimes) | 70 | 19 | 37 | 59 | 39 | 28 |
| Persistence-targeted analysis (across 6 Hr regimes) | 20 | 15 | 6 | 35 | 31 | 19 |

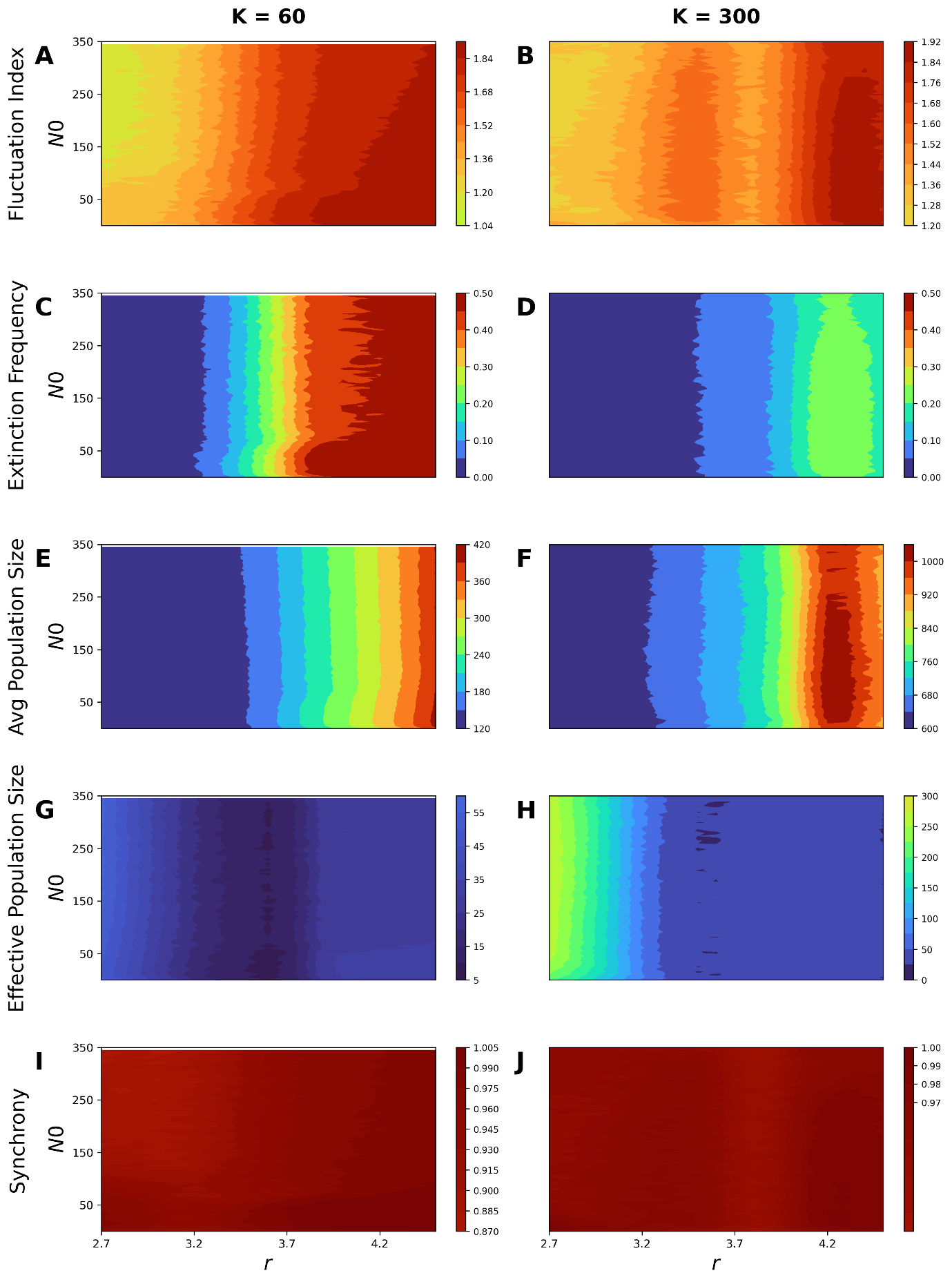

**Figure S1. Sensitivity of stability indices to initial population size (*N₀*).** Heatmaps of (A, B) fluctuation index, (C, D) extinction frequency, (E, F) average population size, (G, H) effective population size, and (I, J) synchrony as functions of N₀ in patch 1 (0–350) and intrinsic growth rate *r* (2.7–4.5), with patch 2 fixed at *N₀* = 20. Left column: *K* = 60; right column: *K* = 300. Dispersal rates: *m₁* = 0.1, *m₂* = 0.3. Colour gradients predominantly follow the *r* axis rather than the *N₀* axis, indicating that stability indices are insensitive to initial population size.

|  | 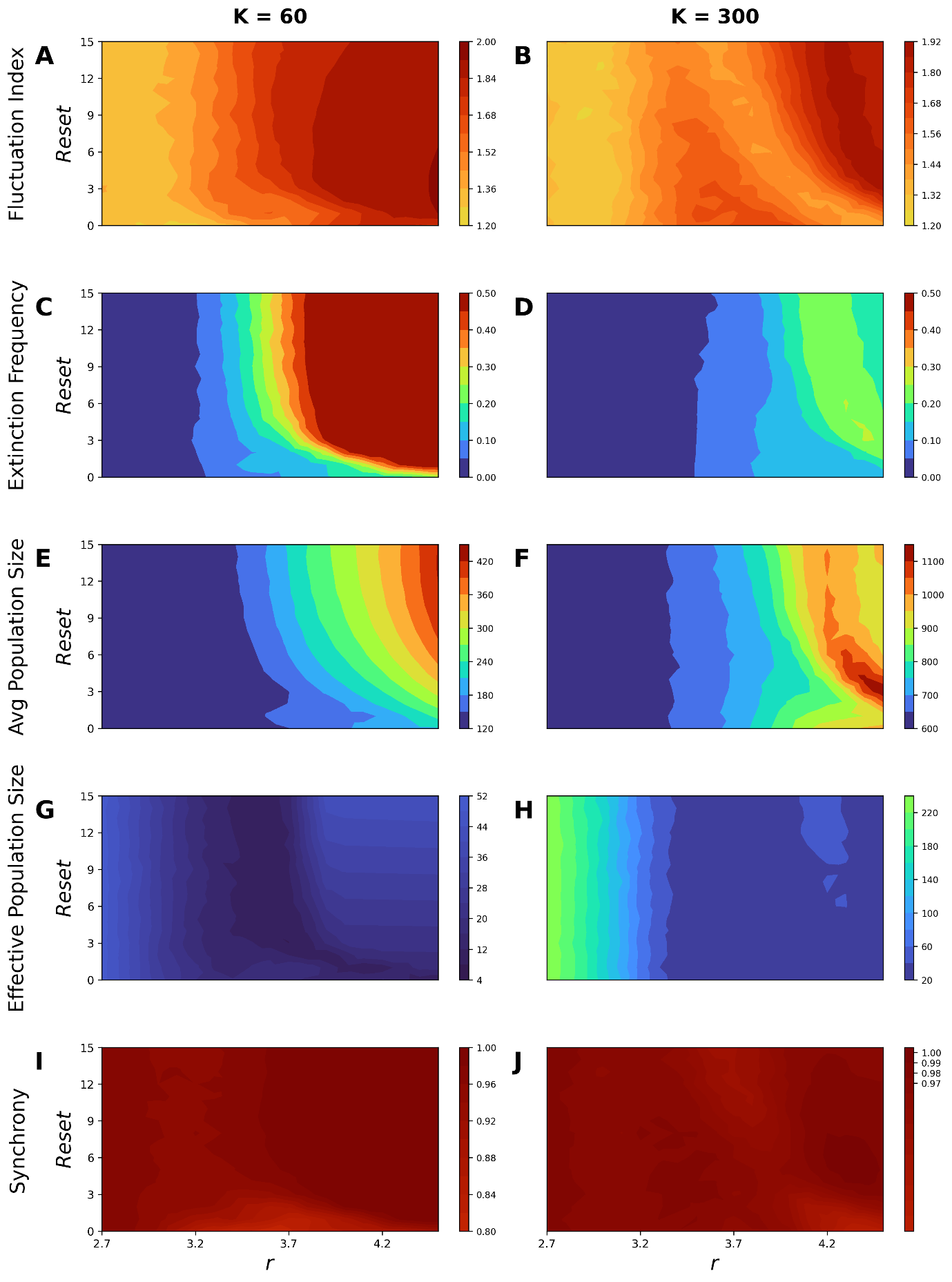 | |
| --- | --- | --- |
| **Figure S2. Sensitivity of stability indices to post-extinction reset value.** Heatmaps of (A, B) fluctuation index, (C, D) extinction frequency, (E, F) average population size, (G, H) effective population size, and (I, J) synchrony as functions of reset value in patch 1 (0–15) and intrinsic growth rate *r* (2.7–4.5), with patch 2 reset fixed at 8. Left column: *K* = 60; right column: *K* = 300. Dispersal rates: *m₁* = 0.1, *m₂* = 0.3. Sensitivity to reset value occurs only when reset ≤ 3 and *r* > 3.7; the default reset value of 8 lies within the insensitive region. | | |
|  | | 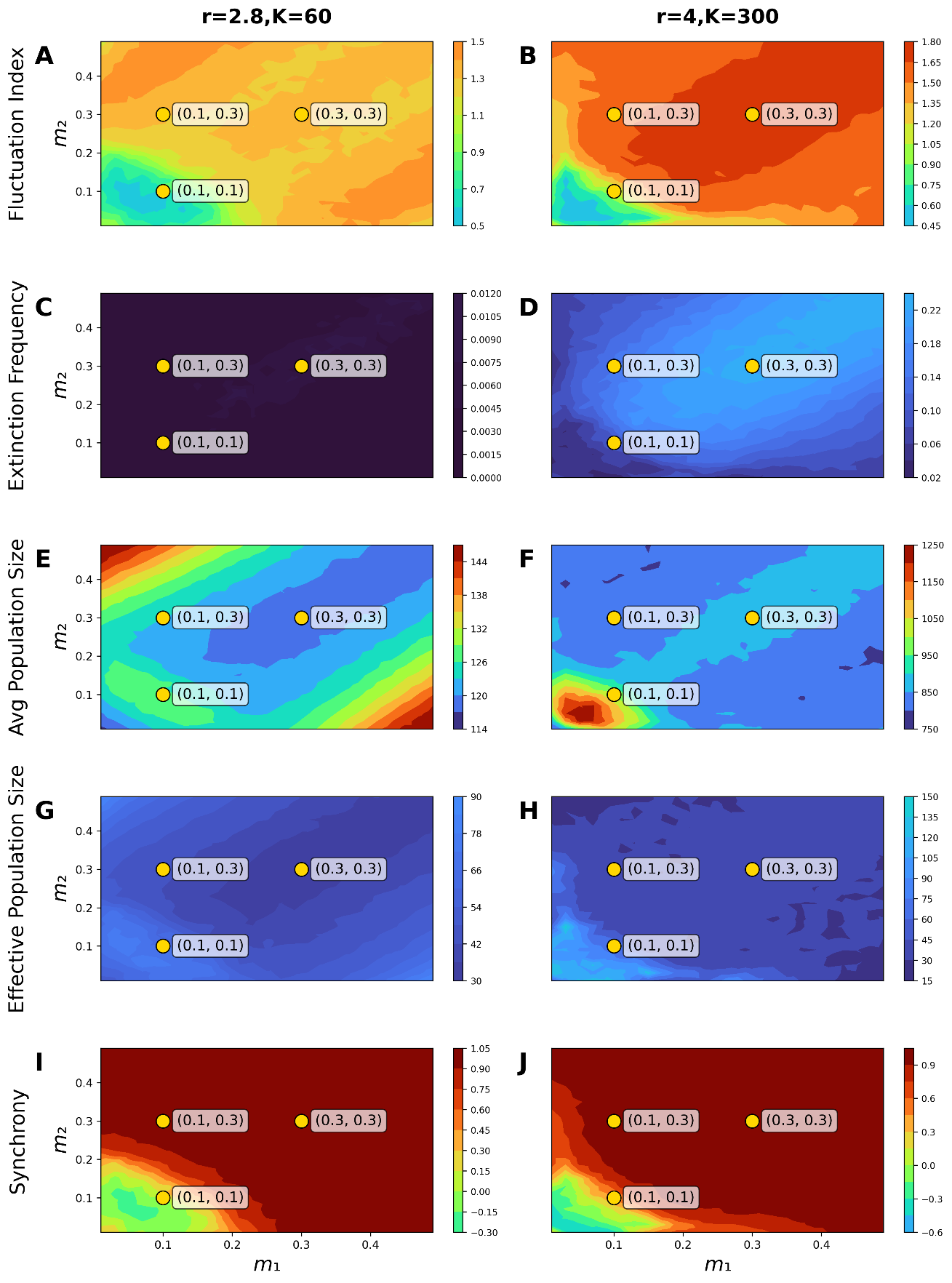 |
|  | | **Figure S3. Effects of asymmetric dispersal on metapopulation dynamics under contrasting intrinsic growth rates and carrying capacities.**  Heatmaps show how five key metrics vary across the two-dimensional dispersal parameter space (m₁, m₂) for two demographic settings: left column (r = 2.8, K = 60) and right column (r = 4.0, K = 300). Warmer colours denote higher values. Panels depict: (A,B) fluctuation index, (C,D) extinction frequency, (E,F) average metapopulation size, (G,H) effective population size, and (I,J) spatial synchrony. Yellow points indicate three focal dispersal combinations (0.1,0.1), (0.1,0.3), and (0.3,0.3) used for detailed analyses in the main text. |

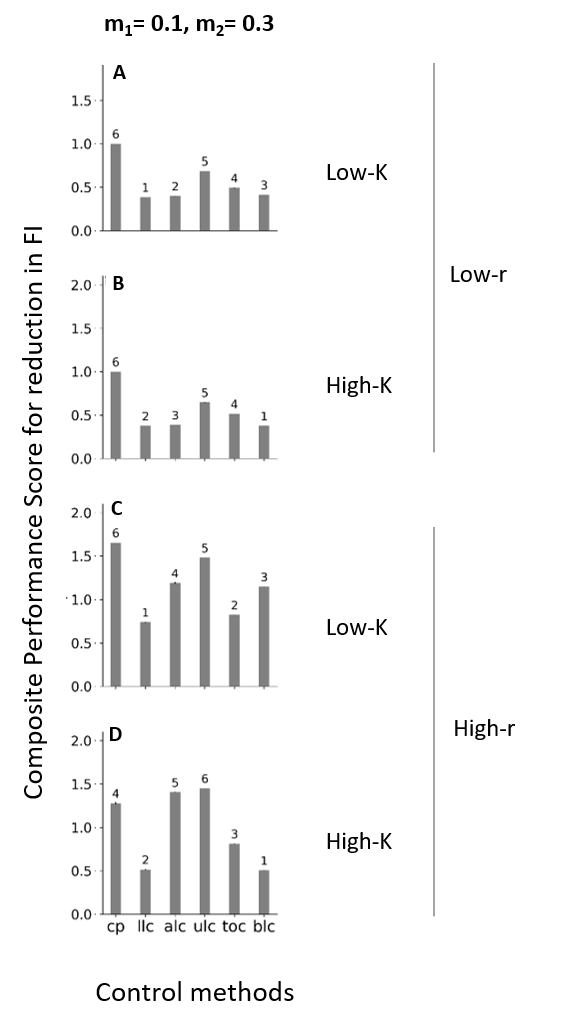

**Figure S4. Composite Performance Scores for 50% Reduction in Fluctuation Index in metapopulations with asymmetric dispersal.** Presented here are the composite performance scores for six population control methods evaluated across four distinct regimes with the objective of achieving a 50% decrease in Fluctuation Index to promote constancy stability. Atop each bar, the numerical annotations represent the performance ranking of the method, with a lower composite score denoting a superior rank. Standard error is depicted as the error bars; however, in most instances, the error bars are too small to be visible.

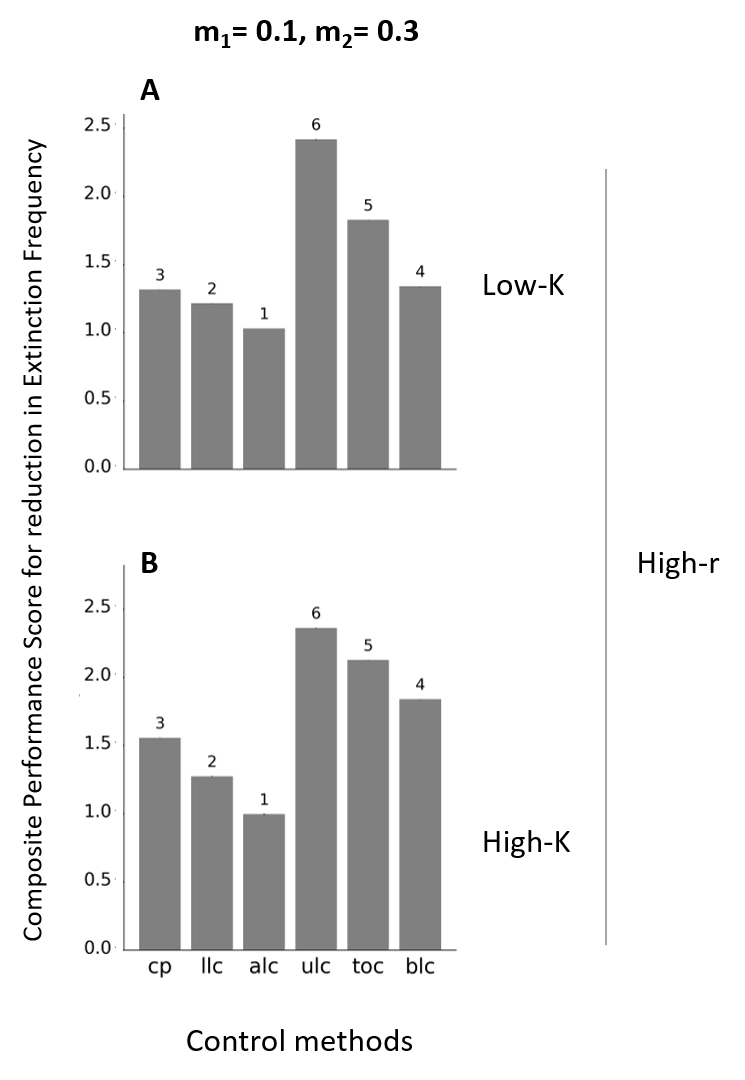

**Figure S5. Composite Performance Scores for 50% Reduction in Extinction Probability in metapopulations with asymmetric dispersal.** Presented here are the composite performance scores for six population control methods evaluated across two regimes, with high intrinsic growth rate (i.e. High-*r* regimes), while achieving a 50% decrease in Extinction Probability to promote persistence stability. Atop each bar, the numerical annotations represent the performance ranking of the method, with a lower composite score denoting a superior rank. Standard error is depicted as the error bars; however, for most instances, the error bars are too small to be visible.

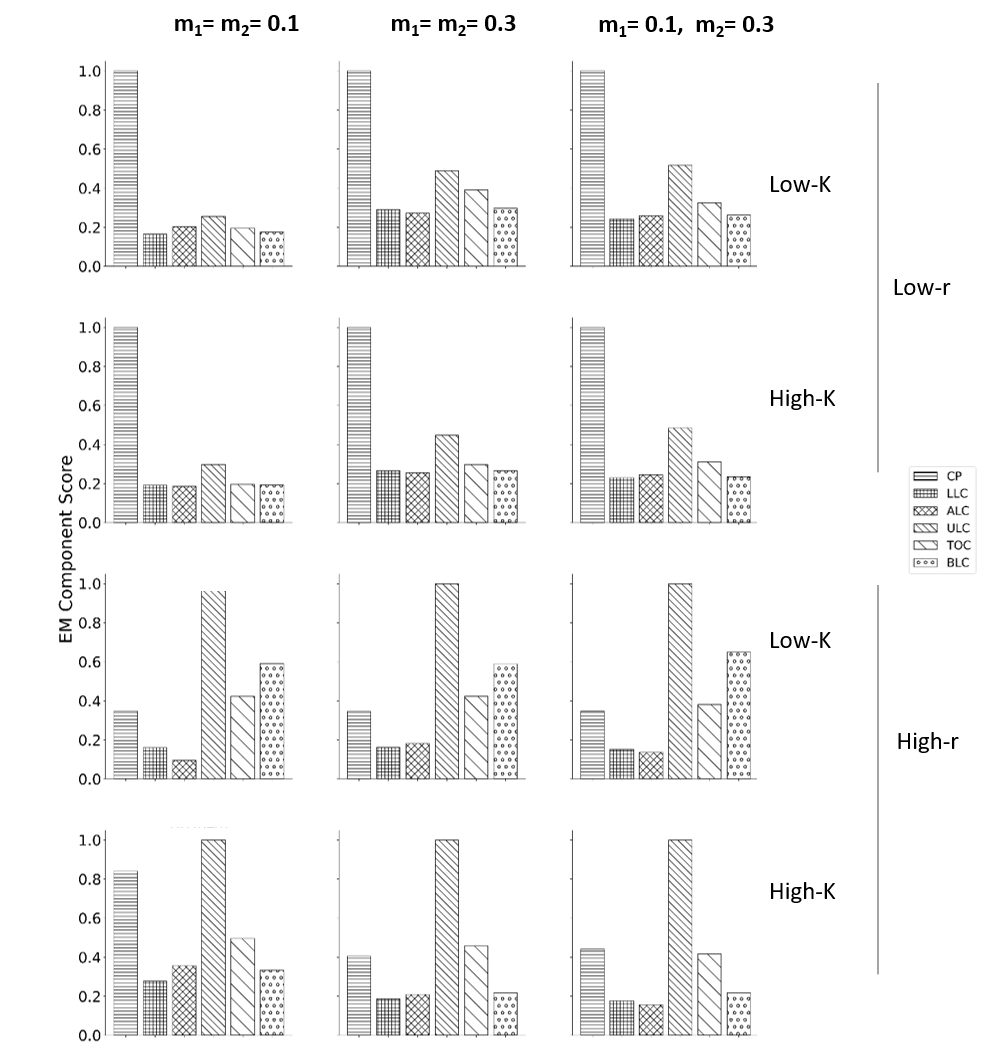

**Fig S6. Component scores for Effort magnitude (*EM*) when reducing *FI* by 50%** employing different control methods across the 12 regimes.

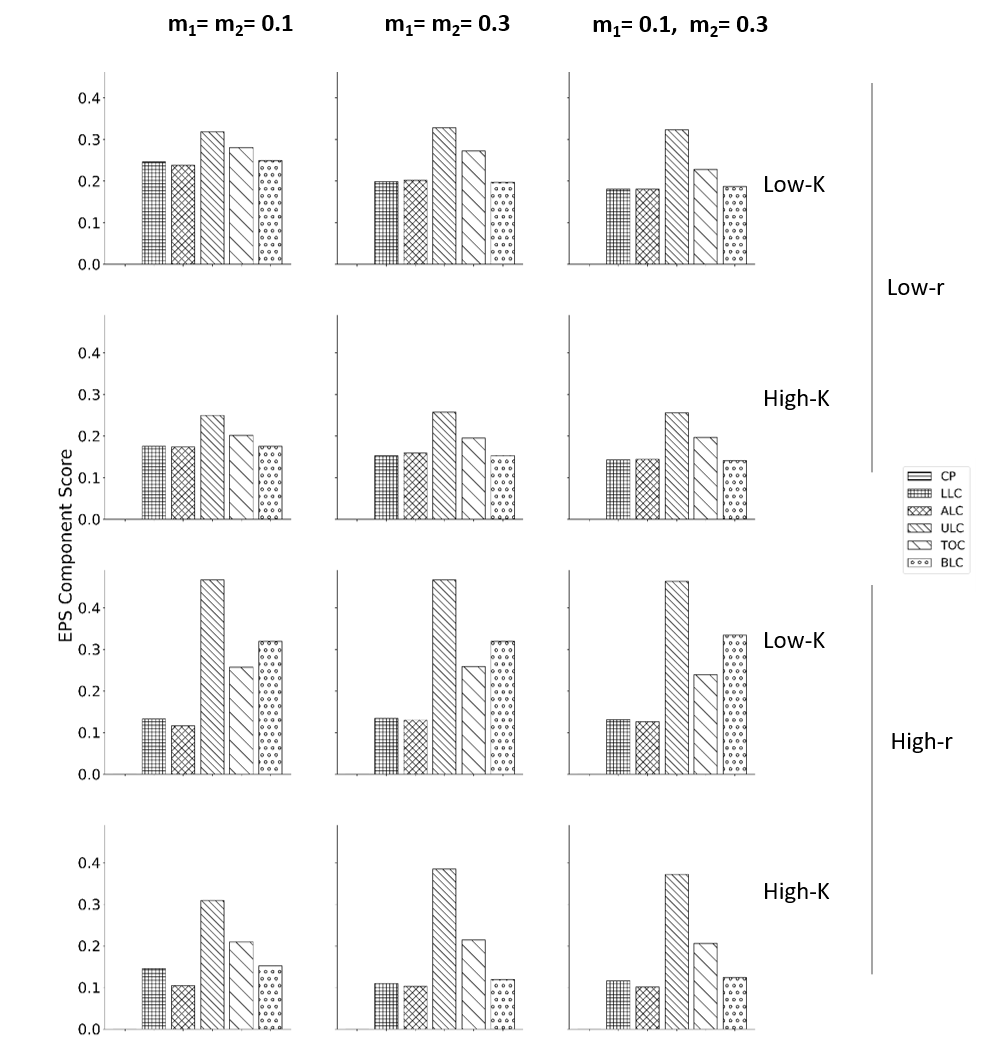

**Fig S7. Component scores for Effective population size (*N_e_*) when reducing FI by 50%** employing different control methods across the 12 regimes.

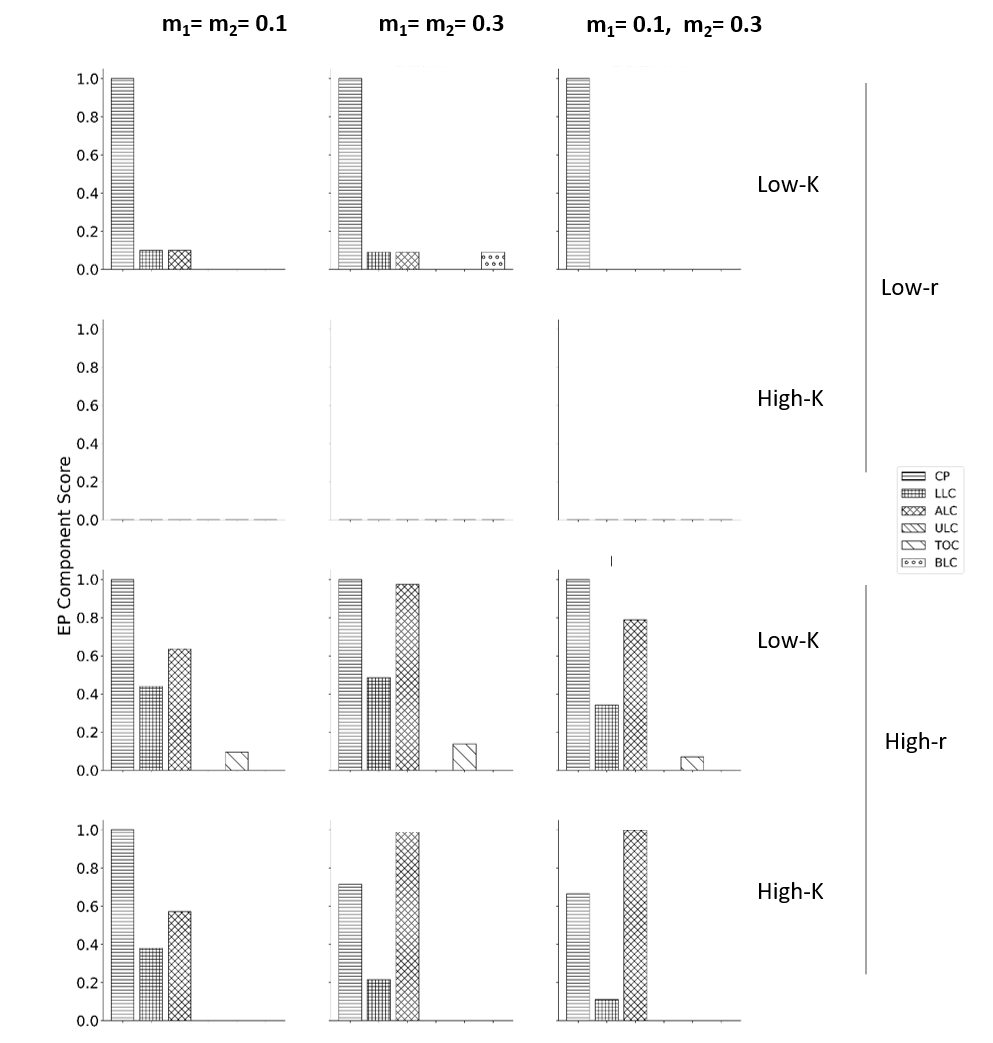

**Fig S8. Component scores for Extinction frequency (*EF*) when reducing *FI* by 50%** employing different control methods across the 12 regimes. The *EF* component index was used in calculating the composite scores only for Hr regimes, since the Lr regimes have negligible *EF*s.

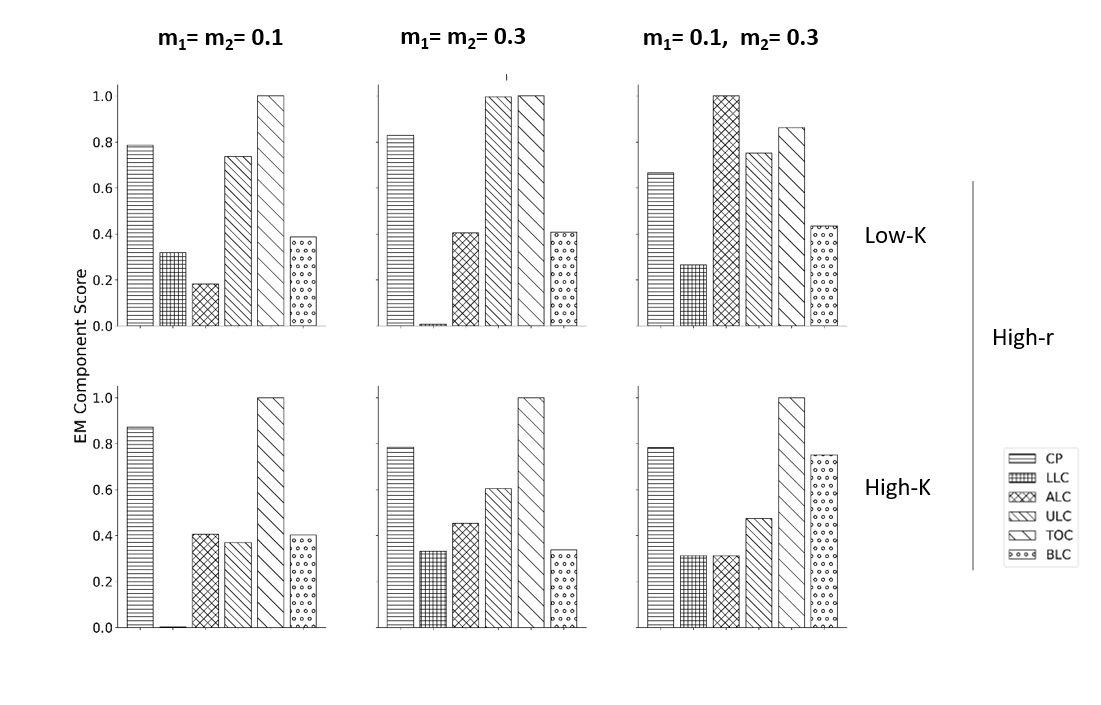

**Fig S9. Component scores for Effort magnitude (*EM*) when reducing extinction frequency by 50%** employing different control methods across the six Hr regimes.

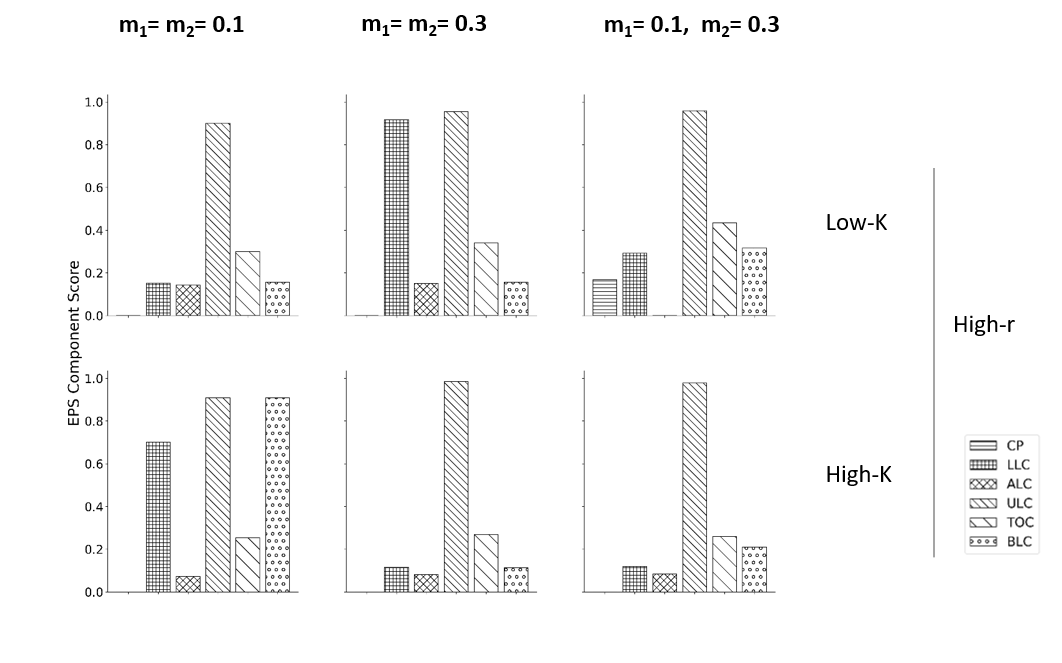

**Fig S10. Component scores for Effective population size (*N_e_*) when reducing extinction frequency by 50%** employing different control methods across the six Hr regimes.

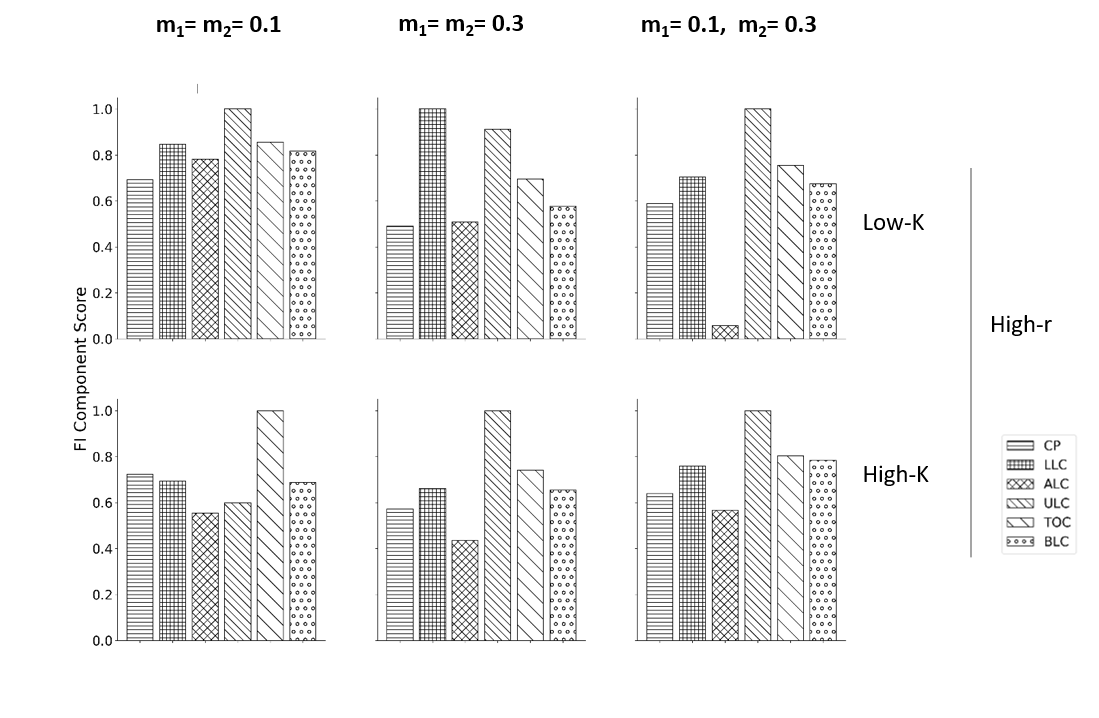

**Fig S11. Component scores for Fluctuation Index (*FI*) when reducing extinction frequency by 50%** employing different control methods across the six Hr regimes.

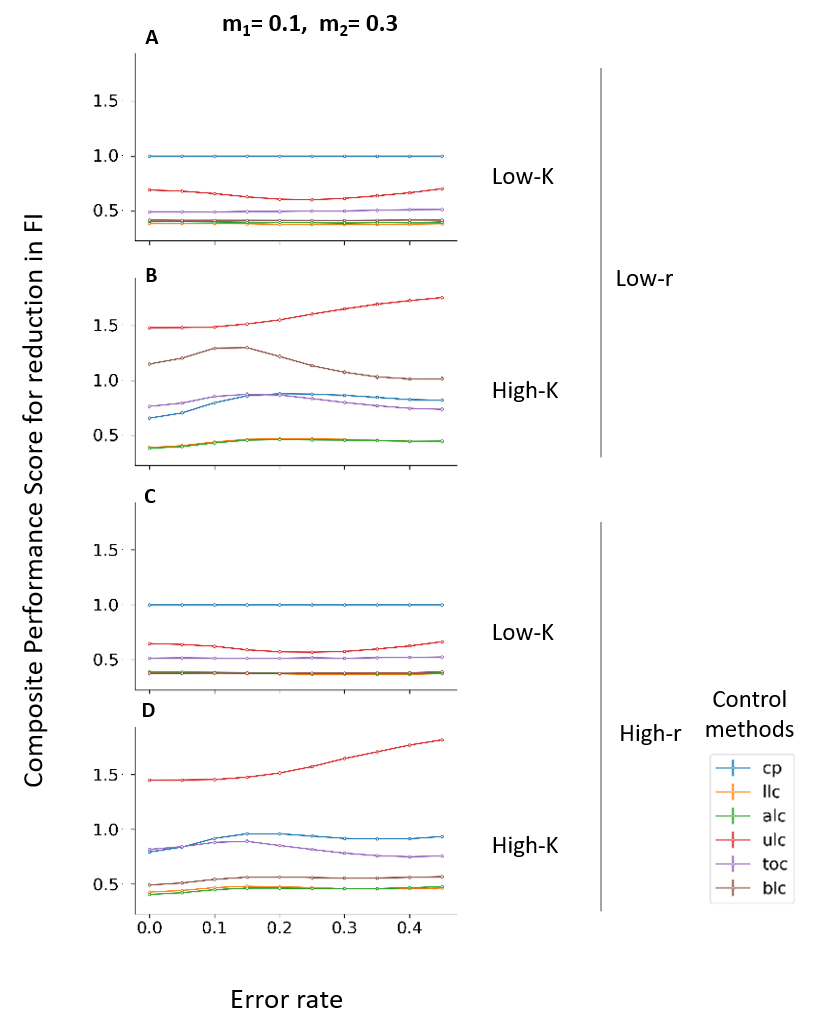

**Fig S12. Change in average composite scores for reducing Fluctuation Index (*FI*) with change in intensity of white noise for regimes with asymmetric dispersal rates.** Corresponding graph for scenarios with symmetric dispersal rates can be found in Figure 5. Note for computing composite performance score, effort magnitude (*EM*) and effective population size (*N_e_*) were used to calculate the composite score for low-r regimes, since they have negligible average extinction frequencies (*EF*) even without any perturbations. For high-*r* regimes, however, Extinction frequency was included in calculating the composite score. The vertical lines on the line plots denote standard error around average composite scores across the replicate runs.

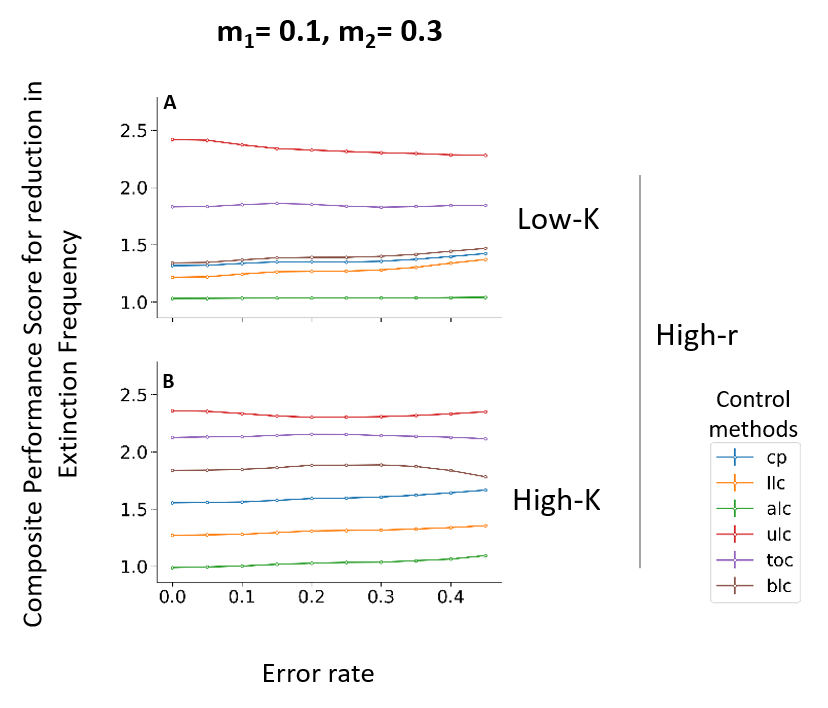

**Figure S13. Change in average composite scores for reducing Extinction Frequency (*EF*) with change in intensity of white noise for regimes with asymmetric dispersal rates.** Corresponding graph for scenarios with asymmetric dispersal rates can be found in supplementary Figure 6. The label on the y axis indicates that Fluctuation index (*FI*), Effort magnitude (*EM*), and Effective population size (*N_e_*) were used to calculate the composite score. The vertical lines on the line plots denote standard error around average composite scores across the replicate runs.

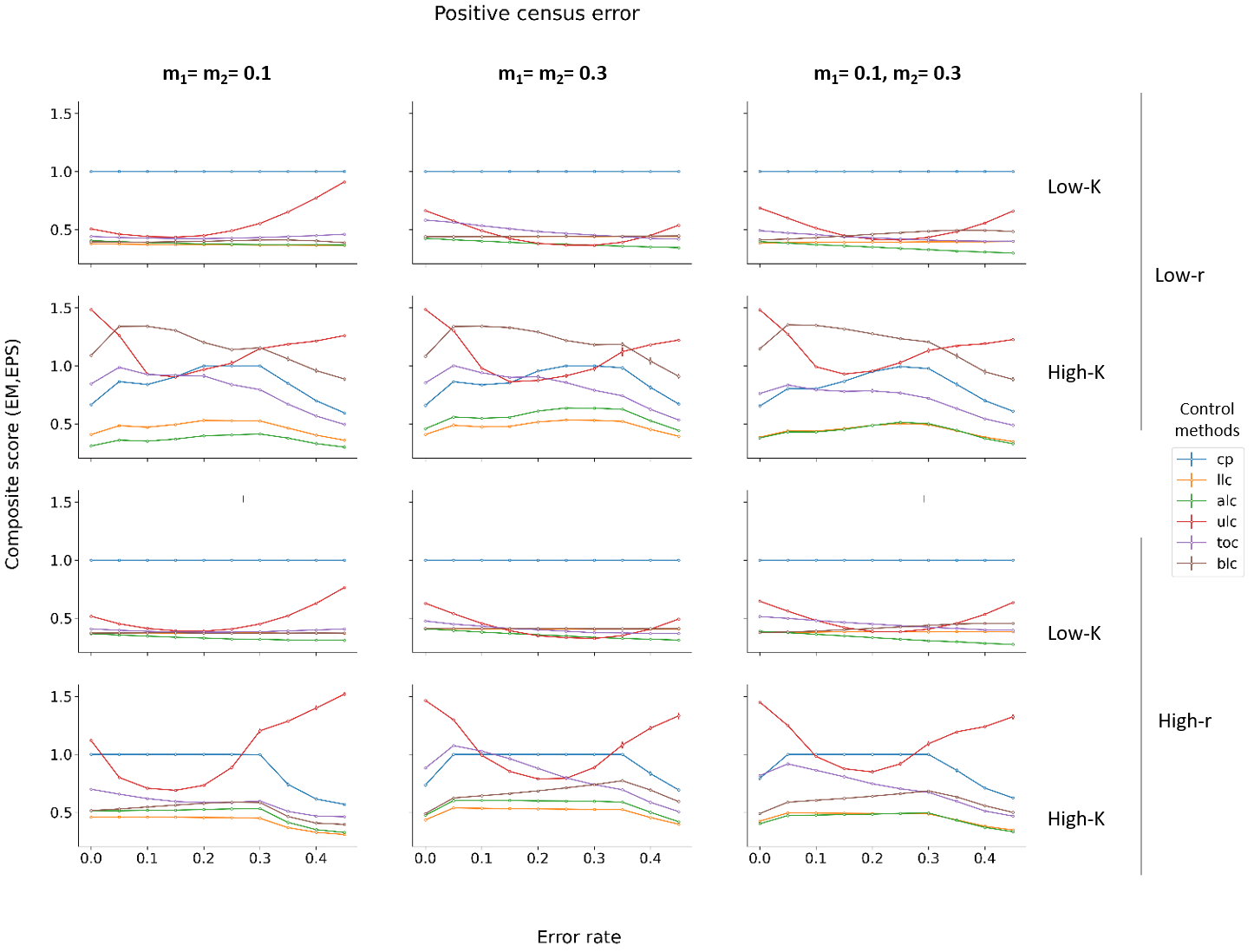

**Fig S14. Change in average composite scores for reducing Fluctuation Index (*FI*) with change in level of positive noise for the 12 regimes.** The label on the y axis indicates that only Effort magnitude (*EM*) and Effective population size (*N_e_*) were used to calculate the composite score for Lr regimes, since they have negligible extinction frequencies (*EF*) even without any perturbations. For Hr regimes, however, Extinction Frequency (*EF*) was also included in calculating the composite score.

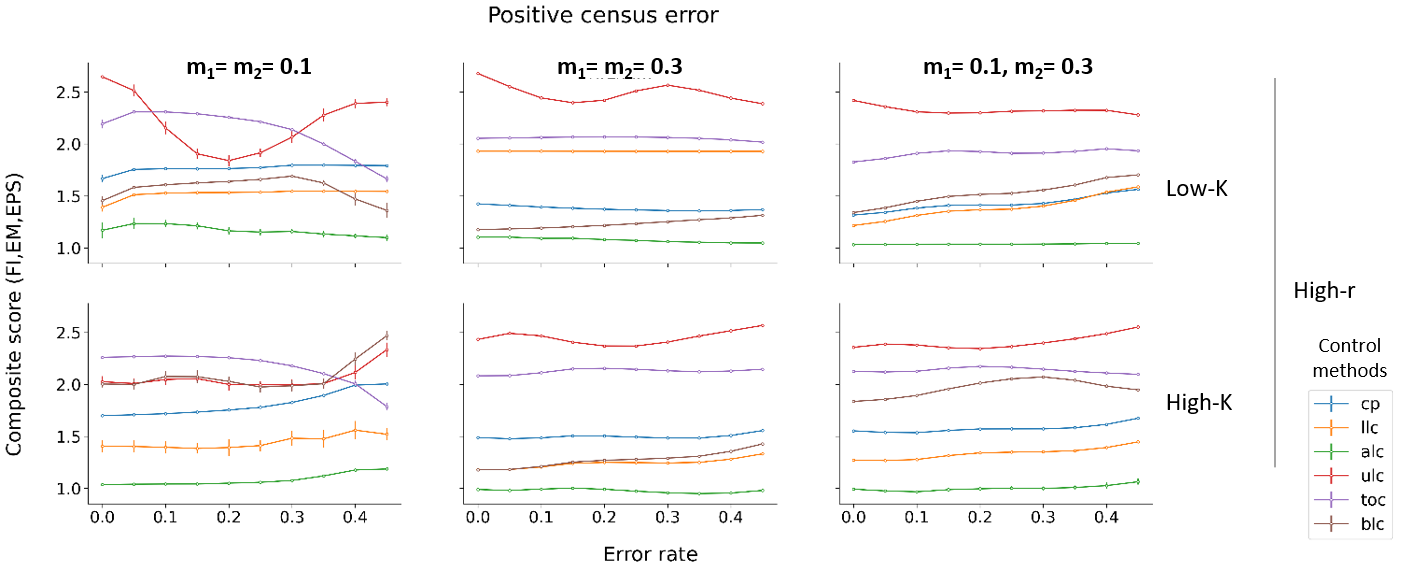

**Fig S15. Change in average composite scores for reducing Extinction Frequency (*EF*) with change in level of positive noise for the 6 regimes.** The label on the y axis indicates that Fluctuation index (*FI*), Effort magnitude (*EM*), and Effective population size (*N_e_*) were used to calculate the composite score.

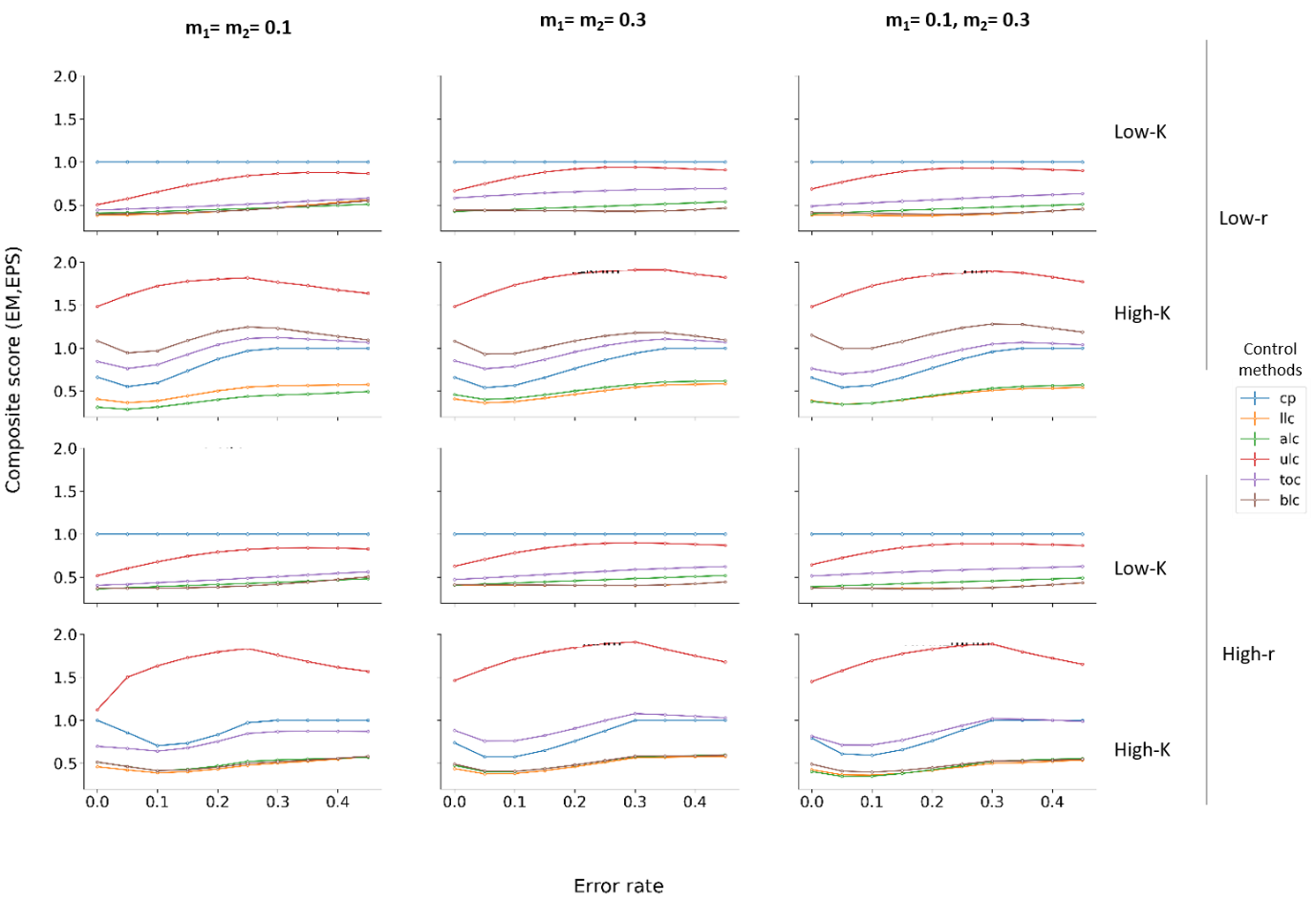

**Fig S16. Change in average composite scores for reducing Fluctuation Index (FI) with change in level of negative noise for the 12 regimes.** The label on the y axis indicates that only Effort magnitude (*EM*) and Effective population size (*N_e_*) were used to calculate the composite score for Lr regimes, since they have negligible extinction frequencies (*EF*). For Hr regimes, however, Extinction Frequency (*EF*), Effort magnitude (*EM*), and Effective population size (*N_e_*), all went into calculating the composite score.

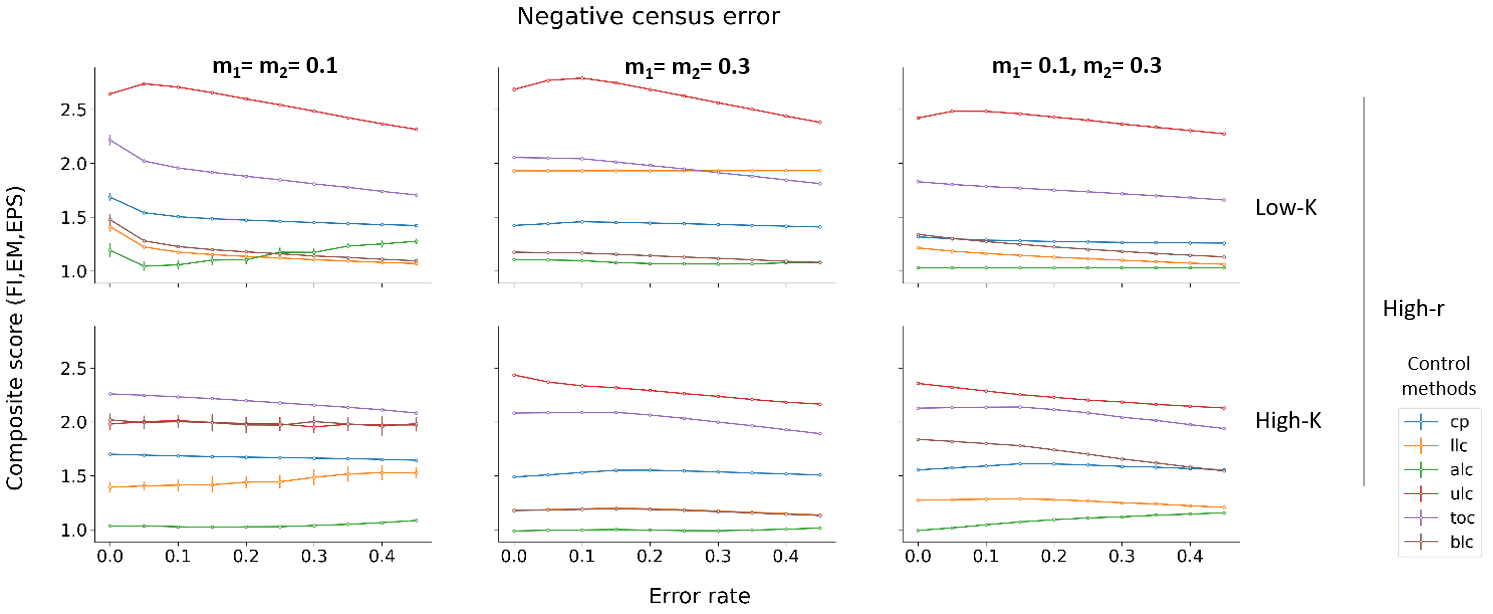

**Fig S17. Change in average composite scores for reducing Extinction Frequency (*EF*) with change in level of negative noise for the 6 regimes.** The label on the y axis indicates that Fluctuation index (*FI*), Effort magnitude (*EM*), and Effective population size (*N_e_*) were used to calculate the composite score.
